# Transcriptomic cytoarchitecture reveals principles of human neocortex organization

**DOI:** 10.1101/2022.11.06.515349

**Authors:** Nikolas L. Jorstad, Jennie Close, Nelson Johansen, Anna Marie Yanny, Eliza R. Barkan, Kyle J. Travaglini, Darren Bertagnolli, Jazmin Campos, Tamara Casper, Kirsten Crichton, Nick Dee, Song-Lin Ding, Emily Gelfand, Jeff Goldy, Daniel Hirschstein, Matthew Kroll, Michael Kunst, Kanan Lathia, Brian Long, Naomi Martin, Delissa McMillen, Trangthanh Pham, Christine Rimorin, Augustin Ruiz, Nadiya Shapovalova, Soraya Shehata, Kimberly Siletti, Saroja Somasundaram, Josef Sulc, Michael Tieu, Amy Torkelson, Herman Tung, Katelyn Ward, Edward M. Callaway, Patrick R. Hof, C. Dirk Keene, Boaz P. Levi, Sten Linnarsson, Partha P. Mitra, Kimberly Smith, Rebecca D. Hodge, Trygve E. Bakken, Ed S. Lein

## Abstract

Variation in cortical cytoarchitecture is the basis for histology-based definition of cortical areas, such as Brodmann areas. Single cell transcriptomics enables higher-resolution characterization of cell types in human cortex, which we used to revisit the idea of the canonical cortical microcircuit and to understand functional areal specialization. Deeply sampled single nucleus RNA-sequencing of eight cortical areas spanning cortical structural variation showed highly consistent cellular makeup for 24 coarse cell subclasses. However, proportions of excitatory neuron subclasses varied strikingly, reflecting differences in intra- and extracortical connectivity across primary sensorimotor and association cortices. Astrocytes and oligodendrocytes also showed differences in laminar organization across areas. Primary visual cortex showed dramatically different organization, including major differences in the ratios of excitatory to inhibitory neurons, expansion of layer 4 excitatory neuron types and specialized inhibitory neurons. Finally, gene expression variation in conserved neuron subclasses predicts differences in synaptic function across areas. Together these results provide a refined cellular and molecular characterization of human cortical cytoarchitecture that reflects functional connectivity and predicts areal specialization.

## Introduction

Cytoarchitectural parcellation of the neocortex has a long history in neuroscience, premised on the idea that structural variations in cellular architecture (*1–3*) and myeloarchitecture (*4*) (representing myelinated fibers or connectivity) underlie functional divisions (reviewed in (*5*)). The neocortex generally has a 6-layered organization that is common across species and areas with a notable exception of agranular areas like the primary motor cortex (M1) that lack an obvious layer 4. Cortical layers contain excitatory projection neurons with generally stereotyped input and output properties that have been hypothesized to represent a “canonical” circuitry (*6, 7*). Despite this common cortical blueprint, cortical areas show relative differences in shape and size, laminar and columnar organization, and proportions of neurons in different layers. Indeed, a century of research has identified an enormous range of cellular properties that vary as a function of positional topography and cortical area (*8–10*), although it has been difficult to gain a clear view on the degree to which there exists a canonical cortical makeup and how reliably to quantify such cytoarchitectural variation.

Single cell and spatial transcriptomic technologies have recently provided new and powerful means to define cortical cellular diversity and spatial organization and compare across cortical areas and species (*11–19*). Applied to human (*11*, *20*), marmoset (*12*), and mouse primary motor cortex (*15*, *16*, *19*), single cell RNA-seq revealed a complex, hierarchical cell type architecture based on gene expression signatures that is quite well conserved across species except at the finest cell type cluster distinctions (*16*). These studies established a subclass level of the hierarchy consisting of 24 well-defined neuronal and nonneuronal types with highly conserved identities across species, distinct laminar patterning and correlated phenotypic properties including cellular anatomy, physiology and broad projection targets (**Table S1**). This cellular architecture provides a comprehensive and discriminatory tool to probe areal variation as well, revealing in mouse cortex that all neuronal subclasses are present across the entire neocortex (*19*), and that areal variation is principally in excitatory neurons rather than inhibitory neurons (*15*). In mouse, monkey and human, the agranular primary motor cortex nevertheless contains layer 4-like neurons despite the lack of a discernable band of these cells characteristic of layer 4 in most other areas (*12*, *21–23*). Single cell RNA-seq, spatial transcriptomics and Patch-seq have revealed a much deeper cellular complexity than subclass in any given cortical area and demonstrated major species differences in relative proportions, cellular properties and cellular microarchitecture (*11*, *12*, *16*, *17*, *24–26*). Much like variation in cortical area devoted to different sensorimotor modalities represents species-specific functional adaptations (*27*), variation in proportions of different cell types likely represent area-specific functional specializations in input-output and local connectivity.

The current study aimed to define quantitative cellular cytoarchitecture across a series of human neocortical areas representative of topographic, functional and structural variation, using deep sampling with single nucleus and spatial transcriptomics methods. The results clearly demonstrate a common canonical architecture of neuronal subclasses, but with wide variation in excitatory neuron subclass proportions that likely reflect different output connectivity and substantial non-canonical areal variation at a finer cell type resolution. Cell type proportions and expression profiles vary as a function of relative proximity on the cortical sheet, with most variation in excitatory neurons but also in inhibitory neurons and astrocytes. The primary visual cortex (V1) was dramatically different from all other cortical areas, with many neuron types only found in V1, reflecting the high degree of specialization for visual processing in the human cortex.

### Within-area cell taxonomies demonstrate common subclass architecture

To sample variation across the cortex, we analyzed eight neocortical areas that include primary (motor, M1; somatosensory, S1; auditory, A1; visual, V1) and association areas (dorsolateral prefrontal cortex, DFC, specifically Brodmann area 9 in the superior frontal gyrus; anterior cingulate cortex, ACC; middle temporal gyrus, MTG; angular gyrus, AnG), span the rostral to caudal (anterior to posterior in many mammals) extent of the cortical sheet, and represent major variations in cortical cytoarchitecture (**Fig. 1A** and **Table S2**) (*28*). Cortical areas were identified across tissue donors using a combination of surface anatomical landmarks and histological verification of cytoarchitecture. Human postmortem brain samples were collected from 5 individuals. Three single nucleus RNA-seq (snRNA-seq) datasets were generated for all areas (except AnG): a 10x Chromium v3 (Cv3) dataset with >924,000 nuclei sampled from all cortical layers, a Cv3 dataset from >231,000 nuclei captured by specific micro-dissection of layer 5 to enrich for rare deep layer excitatory neuron types including layer 5 extratelencephalic projecting (L5 ET) neurons, and a SMART-seqv4 (SSv4) dataset composed of over 60,000 nuclei sampled from individual cortical layers to provide laminar selectivity for all clusters. For AnG, only a Cv3 dataset of all cortical layers was generated (**Fig. 1B**).

**Fig. 1.**
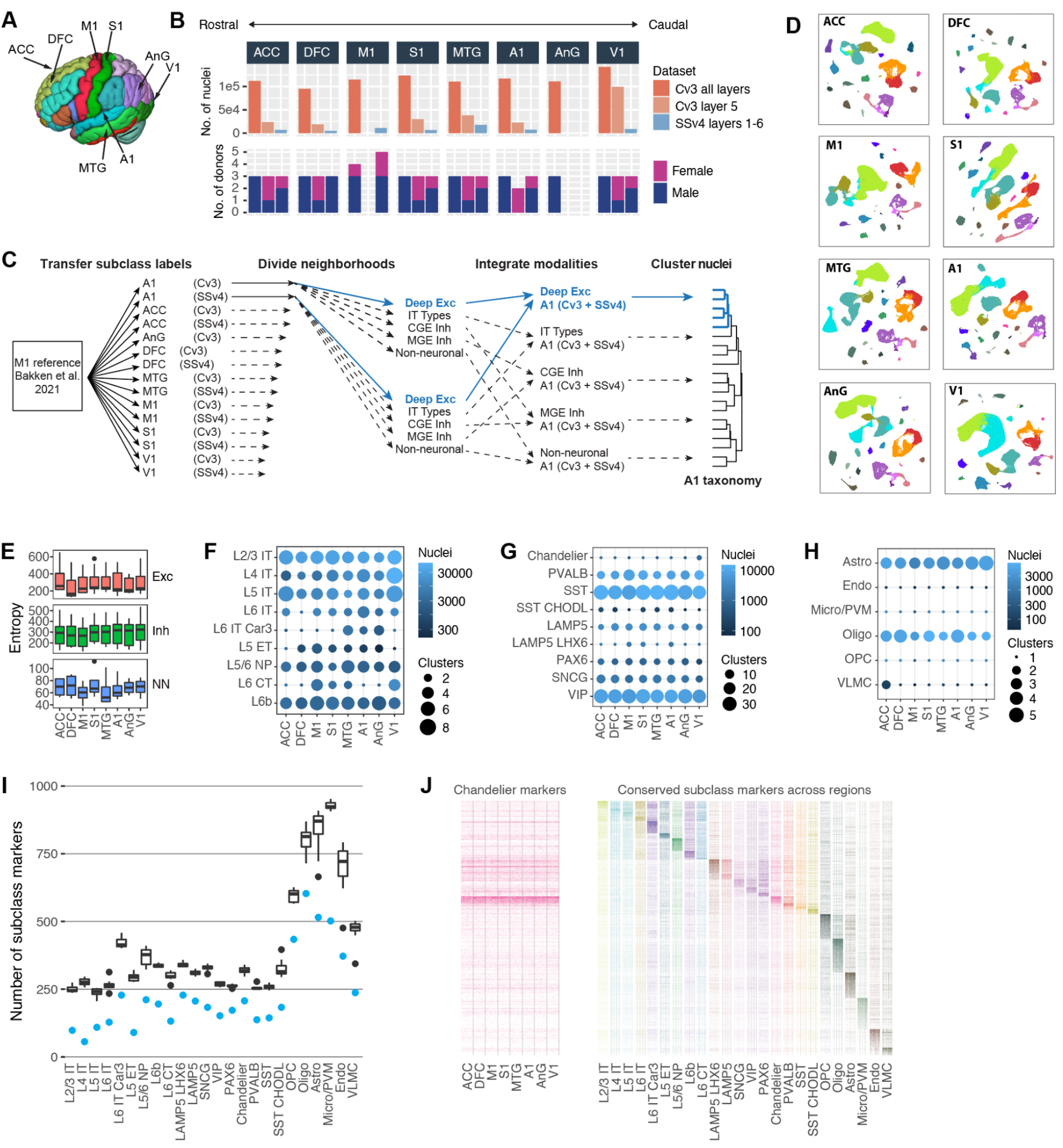
Transcriptomic cell type diversity across human cortical areas. **(A)** Eight areas of the neocortex were sampled from four lobes of the adult human brain. **(B)** snRNA-seq sampling across areas grouped by RNA-seq platform and layer dissection strategy and number of male and female donors. **(C)** Schematic of snRNA-seq clustering to generate cell type taxonomies for each area. **(D)** UMAPs of single nuclei from each area based on variable gene expression and colored by cell subclass as in panel **J**. **(E)** Distributions of subclass transcriptomic entropy are significantly different (P < 0.05) between neuronal (Exc and Inh) and non-neuronal (NN) classes and not between areas based on a two-way ANOVA and post-hoc Tukey HSD tests. **(F,G,H)** Summary of within-area taxonomies showing the number of nuclei sampled from each subclass and the number of distinct clusters (cell types) identified for excitatory (F) and inhibitory (G) neurons and non-neuronal cells (H). **(I)** Number of subclass markers in each area (box plots) and shared across areas (blue points). Box plots show median, interquartile range (IQR), up to 1.5*IQR (whiskers), and outliers (points). **(J)** Heatmaps of conserved marker expression for 50 random nuclei sampled from each area for chandelier interneurons and horizontally compressed for all subclasses.

Nuclei were assigned to one of 24 cell subclasses based on transcriptomic similarity to a reference taxonomy for human M1 (*12, 16*), and subclasses were grouped into five neighborhoods (**Fig. 1C,D**). For each area and neighborhood, nuclei profiled with Cv3 and SSv4 were integrated based on shared coexpression and clustered to identify transcriptomically distinct cell types. Neighborhood clusters were aggregated and organized into within-area taxonomies of between 120 and 142 cell types (**Fig. 1C, S1-S8**) with distinct marker expression (**Table S3**). Cellular variation within subclasses was quantified as the average entropy of variably expressed genes. Entropy was higher for all neuronal than non-neuronal subclasses and not significantly different between excitatory and inhibitory subclasses or across areas based on a two-way ANOVA followed by post-hoc Tukey HSD tests (**Fig. 1E**). Surprisingly, the number of distinct cell types within excitatory subclasses varied across areas (**Fig. 1F**). This within-subclass variation was not driven by differential sampling of nuclei across areas, and was unlike the highly similar diversity of inhibitory neurons and non-neuronal cell types (**Fig. 1G,H**). Of note, there were more layer 5 intratelencephalic projecting (L5 IT) and L4 IT types in V1, the latter expected based on the known expansion and specialization of the thalamorecipient layer 4. L6 IT Car3 neurons were more diverse in MTG, A1, and AnG compared to other areas reflecting a balanced population of *CUX2*-expressing and non-expressing neurons specifically in these cortical areas that we reported as an evolutionary specialization of great apes compared to monkeys (*29*). L5 ET neurons were least diverse in the most rostral area ACC and the most caudal area V1, while layer 6 corticothalamic-projecting (L6 CT) neurons were most diverse in M1, S1, MTG and V1. Individual subclasses had hundreds of distinct markers in each area (**Table S4**), and 20-70% of markers were conserved across areas (**Fig. 1I**). For example, **Fig. 1J** plots expression of a set of chandelier cell markers that were common across areas (left), and a set of common markers for all subclasses (right). Non-neuronal subclasses were transcriptomically distinct and had the most markers, while excitatory subclasses had the smallest fraction of conserved markers, pointing to more variable expression of excitatory neuron gene expression across the cortex as reported in mouse (*15*).

### Cross-areal abundance changes reveal areal specification

The areas analyzed have distinct cytoarchitecture based on conventional Nissl staining that show variation in cell size, shape, laminar and columnar organization (**Fig. 2A**), and spanned the rostrocaudal and mediolateral axes of the cortical sheet (**Fig. 2B**). The relative proportions of transcriptomically defined neuronal subclasses varied across areas (**Fig. 2C** and **Table S5**). Excitatory neuron subclasses showed the greatest differences in proportions across areas that reflect differences in functional architecture. For example, M1 lacks a layer 4 but still has L4 IT neurons at lower proportions than other areas (*12*), and the same was true for agranular ACC, which had the lowest proportion of L4 IT neurons. In contrast, as noted above, V1 has an expanded and highly specialized layer 4 and a corresponding increase in L4 IT neuron proportion. As described previously in mouse cortex (*15*) and between human M1 and MTG areas (*12*), inhibitory neuron subclasses were similar across areas except for a marked increase of the MGE-derived PVALB neurons and fewer CGE-derived interneurons (LAMP5 LHX6, LAMP5, SNCG, VIP, PAX6) in V1. These proportion differences in excitatory and inhibitory neurons were validated *in situ* by labeling neuronal subclasses in MTG and V1 using MERFISH spatial transcriptomics (**Fig. 2C** right panels; **Table S5**), demonstrating they were not an artifact of nuclear isolation and snRNA-seq processing.

**Fig. 2.**
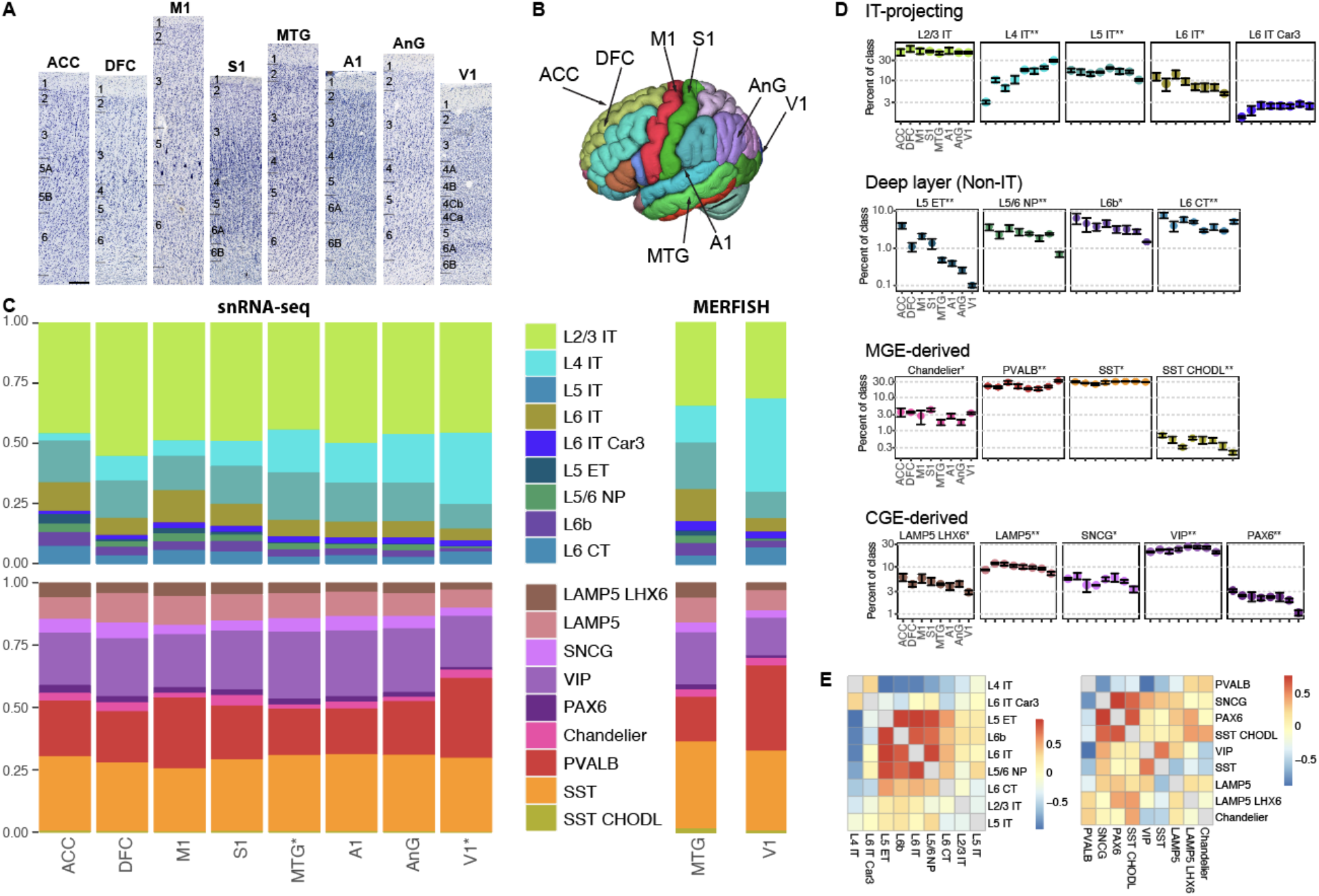
Cell subclass composition reflects cytoarchitecture and varies systematically along the rostrocaudal axis. **(A)** Images of Nissl-stained sections of cortical areas are labeled with approximate layer boundaries and show distinct cytoarchitecture. Areas are ordered by position along the rostrocaudal axis of the cortex. **(B)** Cortical locations of sampled tissue. **(C)** Relative proportions of neuronal subclasses as a fraction of all excitatory or inhibitory neurons in each area and estimated based on snRNA-seq profiling or *in situ* labeling using MERFISH. **(D)** Subclass proportions as a fraction of all neurons in the same class (excitatory or inhibitory) and grouped by neighborhood. Mean +/- standard deviation across donors. Significant differences across regions (ANOVA; *nominal P < 0.05; **Bonferroni-corrected P < 0.05). **(E)** Spearman correlations of excitatory and inhibitory subclass proportions across areas. Scale bar on A, 200 μm.

Subclass proportions were highly consistent across donors (**Fig. 2D**). Examined from a subclass perspective, the most dramatic proportion differences were seen in L4 IT (range 10-fold, from 3-30% of excitatory neurons), and in the much sparser deep subcortically projecting L5 ET neurons (range 50-fold, from 0.1-5%). Unexpectedly, many of these proportion differences varied in a graded fashion generally along the rostrocaudal axis. Pairwise correlations in excitatory neuron proportions revealed correlated rostrocaudal decreases in L5 ET, L6B, L6 IT and layer 5/6 near-projecting neurons (L5/6 NP), with an anticorrelated rostrocaudal increase in L4 IT (**Fig. 2E**). Among inhibitory subclasses, the rarest types (SNCG, PAX6, and SST CHODL) had the most correlated changes in proportions with a decreasing rostrocaudal gradient. PVALB interneurons showed the opposite trend and had increasing proportions in caudal areas, although this trend was variable across areas.

Smaller-scale areal specializations in proportions were overlaid on these broad trends of conservation or rostrocaudal gradients. Notably, many subclasses showed a particularly large difference in V1 (e.g. L5/6 NP, L6B, PAX6). There were more chandelier inhibitory neurons in primary sensory areas (S1, A1, and V1) than expected based on a decreasing rostrocaudal gradient. Also, there were more L4 IT neurons and fewer L5 ET neurons in DFC, more L6 IT neurons in M1, and fewer L6 IT Car3 neurons in ACC than expected based on the broad trends. In summary, cell subclass proportions define a quantitative cytoarchitecture that is canonical in having all 24 subclasses in all areas, with varying proportions and gradient properties that likely reflect developmental gradients with additional specializations driven by the circuit requirements of functionally distinct cortical areas.

### Excitatory to inhibitory neuron ratio varies across cortical areas and layers

In addition to regional specializations in neuronal subclass proportions, we also found regional differences in the relative proportions of excitatory and inhibitory neurons (E:I ratio) (**Table S5**). As previously reported for M1 using snRNA-seq (*12*), we found an E:I ratio of 2:1 was relatively constant across almost all cortical areas, in contrast to the widely reported E:I ratio of 5:1 in mouse. This ratio was much higher in V1 (**Fig. 3A**), with a 4.5:1 ratio comparable to rodents, and we confirmed these values and regional differences with MERFISH analysis (**Table S5**).

**Fig. 3.**
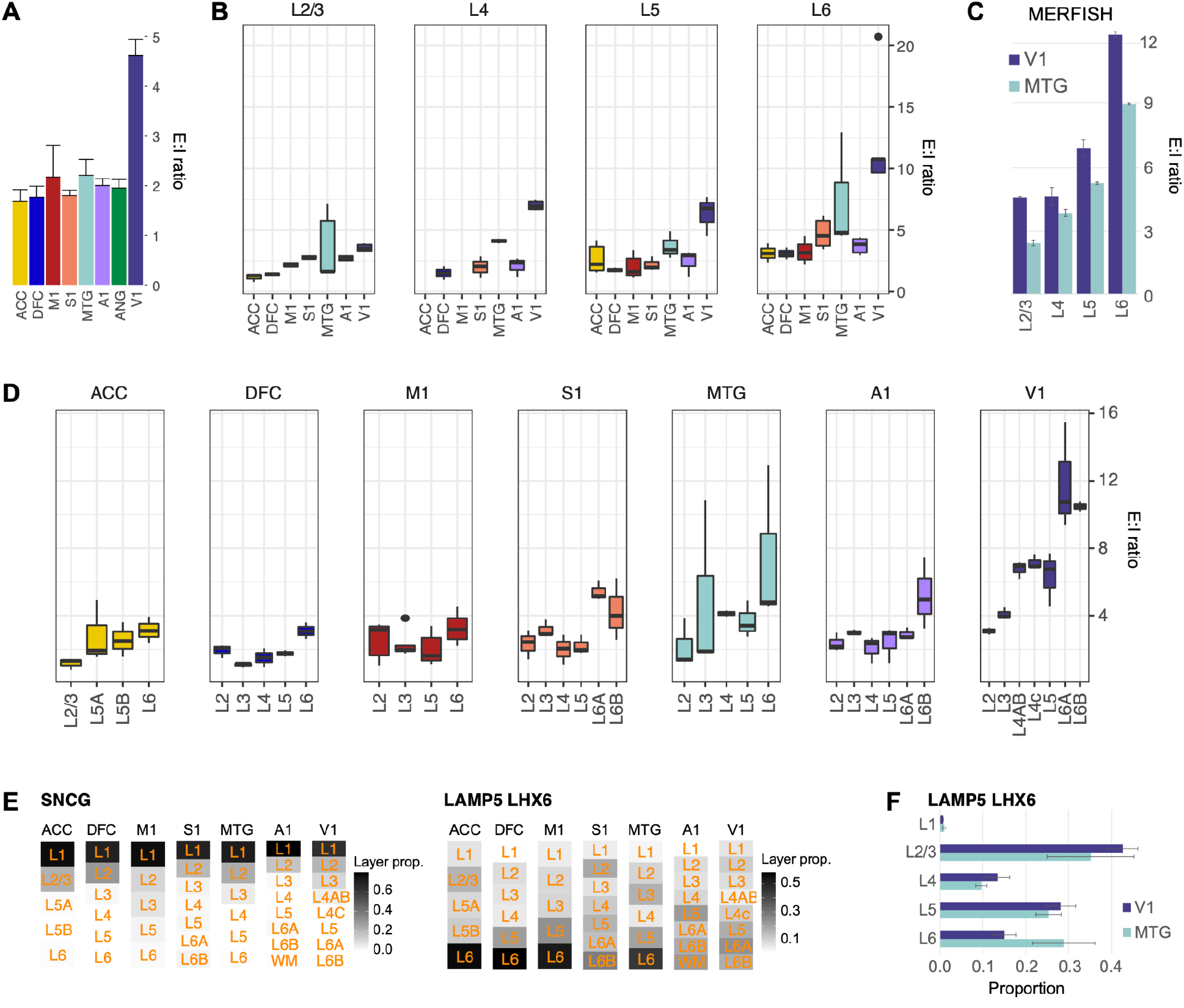
E:I ratio variation across cortical areas and layers. **(A)** Relative number of excitatory neurons to inhibitory neurons (E:I ratio) in each area. Bar plots indicate average and standard deviation across donors. **(B)** E:I ratios estimated for a common set of layers dissected from each area. Box plots show median, interquartile range (IQR), up to 1.5*IQR (whiskers), and outliers (points) across multiple donors. **(C)** Validation of increased E:I ratios in all cortical layers in V1 compared to MTG based on MERFISH experiments. Bar plots and whiskers indicate average and standard deviation of E:I ratios across donors, respectively. **(D)** E:I ratios estimated for all layers dissected from each area. **(E)** Laminar distributions of interneurons were conserved (SNCG) or divergent (LAMP5 LHX6) across areas based on counts of layer-dissected nuclei. Note that primary sensory areas (S1, A1 and V1) have a distinct distribution of LAMP5 LHX6 neurons. **(F)** MERFISH *in situ* labeling of LAMP5 LHX6 cells shows a decreased proportion of cells in layer 6 of V1 compared to MTG.

The layer-specific dissections for nuclei profiled with SSv4 from 7 regions (all except AnG) allowed a deeper exploration of E:I ratio variation, and demonstrated that a single ratio is insufficient to describe that variation. E:I ratios varied by area and layer, and were highly consistent across donors (**Fig. 3B**). Increased variability was seen in MTG that may be due to inconsistent sampling of different subregions of this large cortical area. V1 had the highest ratio in all layers, not just in layer 4 (7:1) but also in layers 5 and 6, where the highest E:I ratio of 10:1 was seen. Moreover, there was a monotonic increase in the E:I ratio of the other areas along a rostrocaudal gradient, and this was most apparent in L2/3. E:I ratios were more variable in layers 4 and 5, masking the trend in overall E:I ratios (**Fig. 3A**). Laminar *in situ* counts of excitatory and inhibitory neurons in MTG and V1 using MERFISH confirmed a consistently higher E:I ratio in all layers of V1 (**Fig. 3C**). From a within-area perspective, the E:I ratios consistently increased with cortical depth, with the highest ratios in layer 6 for all areas (**Fig. 3D**). Furthermore, the E:I ratio in layer 4 was uniquely elevated in V1 relative to layers 2, 3 and 5, highlighting a further specialization of visual processing compared to other sensory modalities. Finally, laminar distributions of excitatory and inhibitory neurons were more consistent across cortical areas (**Fig. S9**), such as SNCG in layer 1 (**Fig. 3E**). Some areal and laminar variation was seen as well, such as LAMP5 LHX6 proportions in layer 6 that was seen with RNA-seq and confirmed with MERFISH (**Fig. 3F**). Taken together, E:I ratios vary extensively both by layer and area, with dramatically different ratios in V1 and areal variation that is masked by averaging across cortical layers.

### Transcriptomic cellular topography

To characterize the transcriptomic landscape of neuronal subclasses across the cortex, neuronal single nuclei were integrated by donor for each of four neighborhoods (IT-projecting glutamatergic, Deep layer (Non-IT) excitatory, MGE-derived GABAergic and CGE-derived GABAergic) and visualized as UMAPs colored by subclass (**Fig. 4A**) and cortical area (**Fig. 4B**). Three clear organizational principles were apparent. First, excitatory neurons had strong areal signatures, seen by the clear banding by area, while inhibitory neurons were almost completely intermixed across areas similar to reports in mouse cortex (*15*). Second, there were dramatic specializations in V1. For example, there was a massive expansion of L4 IT neurons, and V1 appeared as separate islands for most IT-projecting subclasses and particularly for L6 CT neurons. Distinct V1 islands were also seen for parts of the PVALB and SST subclasses (arrows in MGE-derived UMAPs). Third, the areal similarity of excitatory neurons appeared to vary in a rostrocaudal topographic order for many subclasses, similar to prior reports of gene expression similarity across human cortex (*30*). Neighboring areas were extremely similar and intermixed despite functional distinctiveness; for example, nuclei from M1 and S1 were extensively intermingled despite clear functional specificity for motor and somatosensory functions, respectively.

**Fig. 4.**
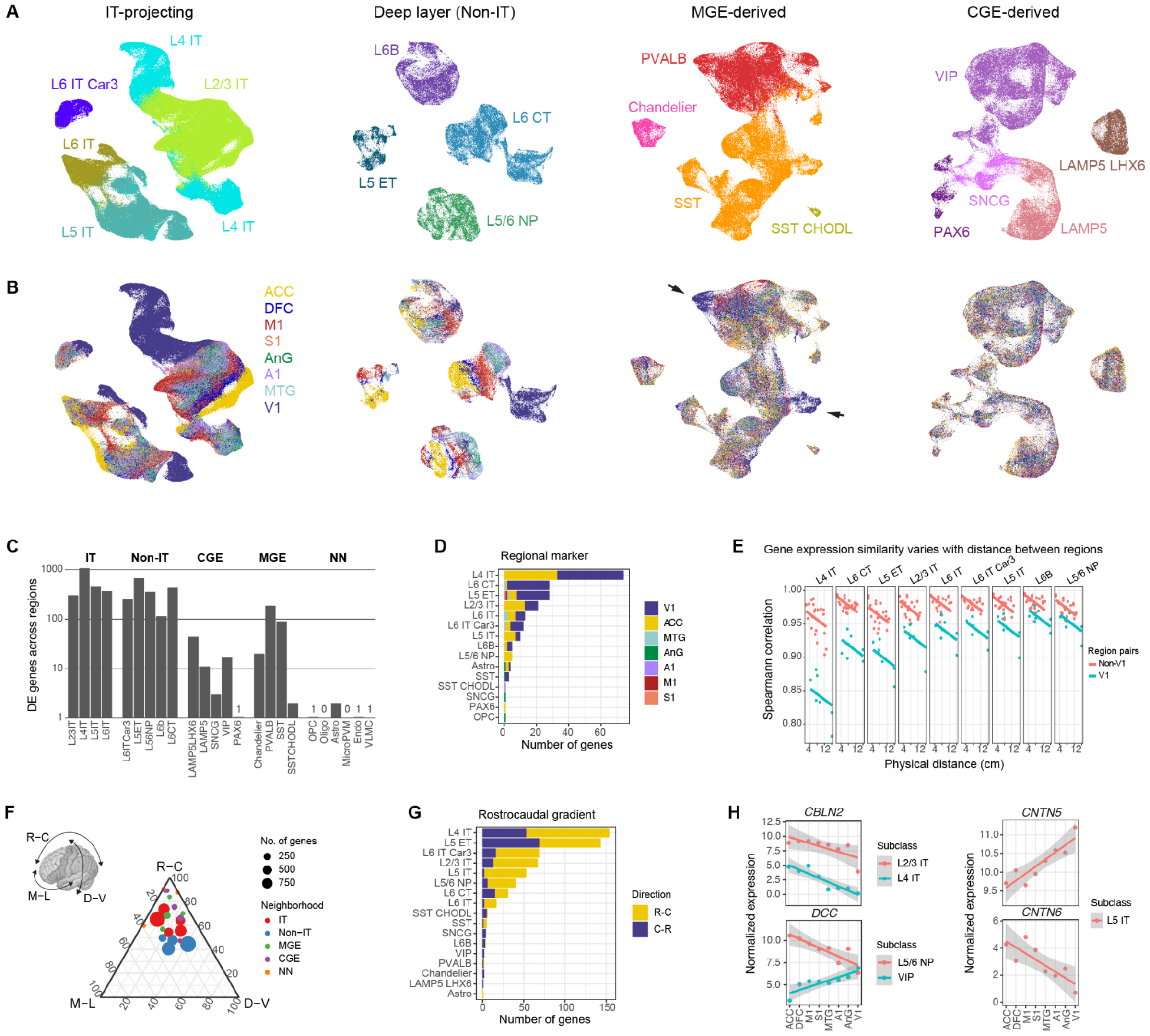
Transcriptional topography across cortical areas. **(A,B)** UMAPs showing transcriptomic similarities of single nuclei dissected from eight cortical areas and colored by neuronal subclass (A) and area (B) for excitatory and inhibitory neuron neighborhoods. Arrows mark V1-specialized PVALB and SST neurons. **(C)** The number of genes that are significantly differentially expressed across areas for each subclass grouped by neighborhood (ANOVA, Bonferroni adjusted P-value < 0.05). Subclasses with 0 or 1 DEG are labeled. See **Table S6** for all DEGs. **(D)** The number of genes that have highly enriched expression in a single area for each subclass. **(E)** Spearman correlations of expression similarity between pairs of areas as a function of the approximate physical distance along an unfolded neocortical sheet. Pairwise comparisons that include V1 (blue points) or do not include V1 (red) are grouped separately because V1 is so transcriptomically distinct. Intercepts but not slopes are significantly different across subclasses based on a linear model. **(F)** Ternary plot summarizing the relative proportion of variance explained by expression gradients across areas along rostrocaudal (R-C), mediolateral (M-L), and dorsoventral (D-V) axes for each subclass. Point size indicates the number of genes with >5% of variance explained by at least one gradient, and point location shows the weighted mean proportion across all genes (shown in **Fig. S10H**). Points are colored by cell neighborhood, and most cells have predominantly R-C gradients, while deep layer (non-IT) neurons have more variable expression across the cortex (blue points are near the center of the plot). **(G)** For each subclass, the number of genes with expression that increases (R-C) or decreases (C-R) in areas ordered by their position along the rostrocaudal axis. **(H)** Examples of genes with rostrocaudal gradient expression that have been previously described in development (*CBLN2*) (*32*), has opposing gradients in different subclasses for the same gene (*DCC*) or for two related genes (*CNTN5* and *CNTN6*) involved in neuronal connectivity for the same subclass.

Areal variation in gene expression mirrored the UMAP trends. The number of differentially expressed genes (DEGs, **Table S5**) across areas was largest for excitatory neurons (**Fig. 4C**), with the highest numbers for L4 IT and L5 ET neurons (over 1000 DEGs). DEGs for inhibitory neuron subclasses varied widely, from over 100 DEGs for SST and PVALB interneurons to fewer than 10 DEGs for SNCG and SST CHODL and a single DEG (*ADAMTS9-AS2*) for PAX6. Non-neuronal cell subclasses similarly displayed few areal DEGs. We next identified area-specific markers (**Table S7**), which were much less common, using a previously defined tau score (*31*). Excitatory neurons expressed the vast majority of these highly specific areal markers, and ACC and V1 were the most unique areas (**Figs. 4D, S10A,B**). IT-projecting neurons were specialized in both ACC and V1, while non-IT L6 CT and L5 ET neurons were specialized mostly in V1. In other words, the vast majority of area-specific gene markers were found in the rostral most (ACC) and caudal most (V1) areas.

The topographic ordering of the excitatory neuron subclasses above suggested systematic graded changes as a function of distance, similar to bulk tissue profiling studies reporting gradual changes in gene expression across the cortical sheet (*30*). We therefore calculated transcriptomic similarities of excitatory subclasses as a function of the approximate physical distance between pairs of areas on an unfolded cortical sheet (**Fig. 4E** and **Table S2**). Because V1 was so transcriptomically distinct (**Fig. 4A**), we fit two linear models of subclass similarity versus areal distance, one that included pairwise comparisons to V1 and one that did not (**Fig. 4E**). Strikingly, all excitatory neuron subclasses showed the same monotonic decrease of similarity with distance, but had different amounts of transcriptomic specialization in V1 (i.e. intercepts but not slopes are significantly different in **Fig. 4E**). Interneuron similarity also decreased with distance at the same rate for all subclasses, albeit at about 40% the rate of excitatory neurons, and with much less specialization in V1 (**Fig. S10C**). In contrast, non-neuronal expression did not change systematically with inter-areal distance and was not more specialized in V1 (**Fig. S10D**).

To determine more precisely how gene expression varied across the cortical sheet, we performed a variance partitioning analysis for each subclass (**Fig. S10E** and **Table S8**). More genes were explained by area identity and position along spatial gradients for excitatory than inhibitory neurons or non-neuronal cells as observed in the UMAPs, although the proportion of variance explained was comparable (**Fig. S10F**). Among IT-projecting neurons, some genes showed unique patterning in a single subclass while other genes were topographically patterned in all IT subclasses (**Fig. S10G**). We calculated the expression variance explained by gradients along three axes: rostrocaudal (R-C), mediolateral (M-L), and dorsoventral (D-V). For genes with at least 5% of expression variance explained by any gradient, we then quantified the relative strength of gradients based on the relative proportion of expression variance that was explained (**Fig. S10H**). For most subclasses, rostrocaudal gradients were dominant, except Non-IT subclasses also expressed many genes with mediolateral and dorsoventral gradients (**Fig. 4F, S10H**).

We robustly defined a set of rostrocaudal genes for each subclass by requiring a Spearman correlation >0.7 between expression and areal position along the rostrocaudal axis and a correlation >0.5 after excluding V1 and ACC. For the most varying L4 IT and L5 ET neurons, roughly equal numbers of genes increased and decreased expression rostrocaudally (**Fig. 4G**). In contrast, for other subclasses, many more genes increased rather than decreased expression along the rostrocaudal axis. For most neuronal subclasses, the correlations of rostrocaudal genes were greater than correlations to a randomly shuffled ordering of areas (**Fig. S10I**). Genes with a rostrocaudal gradient in one subclass tended to have a gradient in the same direction in other subclasses that expressed the gene (**Fig. S10J**), such as *CBLN2* in L2/3 IT and L4 IT neurons (**Fig. 4H**), which is expressed in a similar gradient in maturing cortical neurons during human prenatal development (*32*). However, some genes such as *DCC* had opposing gradients in different subclasses (L5/6 NP and VIP), and some functionally related genes had opposing gradients in the same subclass, such as the cell adhesion molecules Contactin 5 and 6 (*CNTN5* and *CNTN6*) in L5 IT neurons (**Fig. 4H**). Based on Gene Ontology (GO) analysis, genes with strong areal enrichment or rostrocaudal gradients included voltage-gated potassium and calcium channels. Interestingly, only rostrocaudal genes were associated with axon guidance pathways including SLIT/ROBO, ephrin, and semaphorin signaling molecules (**Table S7**) that likely reflects developmental patterning of connectivity.

### Cross-areal consensus taxonomy

We next moved from the relatively coarse level of cell subclasses to understand areal variation at the finer level of cell types, by clustering the integrated cell neighborhoods shown in **Figure 4** to identify a set of cell types either common to or varying across cortical areas (**Fig. 5A**). We defined and organized 153 cell types by transcriptomic similarity into a consensus taxonomy (**Fig. 5B**). Consensus cell types had consistent markers across areas (**Table S9**), were represented in all donors (**Fig. 5C**) and ranged from 0.01% to 20% of excitatory and inhibitory neurons and from 0.1% to 30% of non-neuronal cells (**Fig. 5D**). The majority of types were found in all eight areas, with particularly uniform representation across areas for most inhibitory neuron types and non-neuronal cells (**Fig. 5E**). However, there were clearly area-enriched or area-specific cell types, most notably in V1 (dark blue). V1-enriched clusters were seen in nearly all excitatory subclasses, particularly L4 IT, as well as SST, and a few PVALB and VIP types. There was also one ACC-selective VIP type. Other notable patterns included the frequent cross-areal excitatory cell types common to M1 and neighboring S1, or nearby MTG and A1, again reflecting a similarity by proximity.

**Fig. 5.**
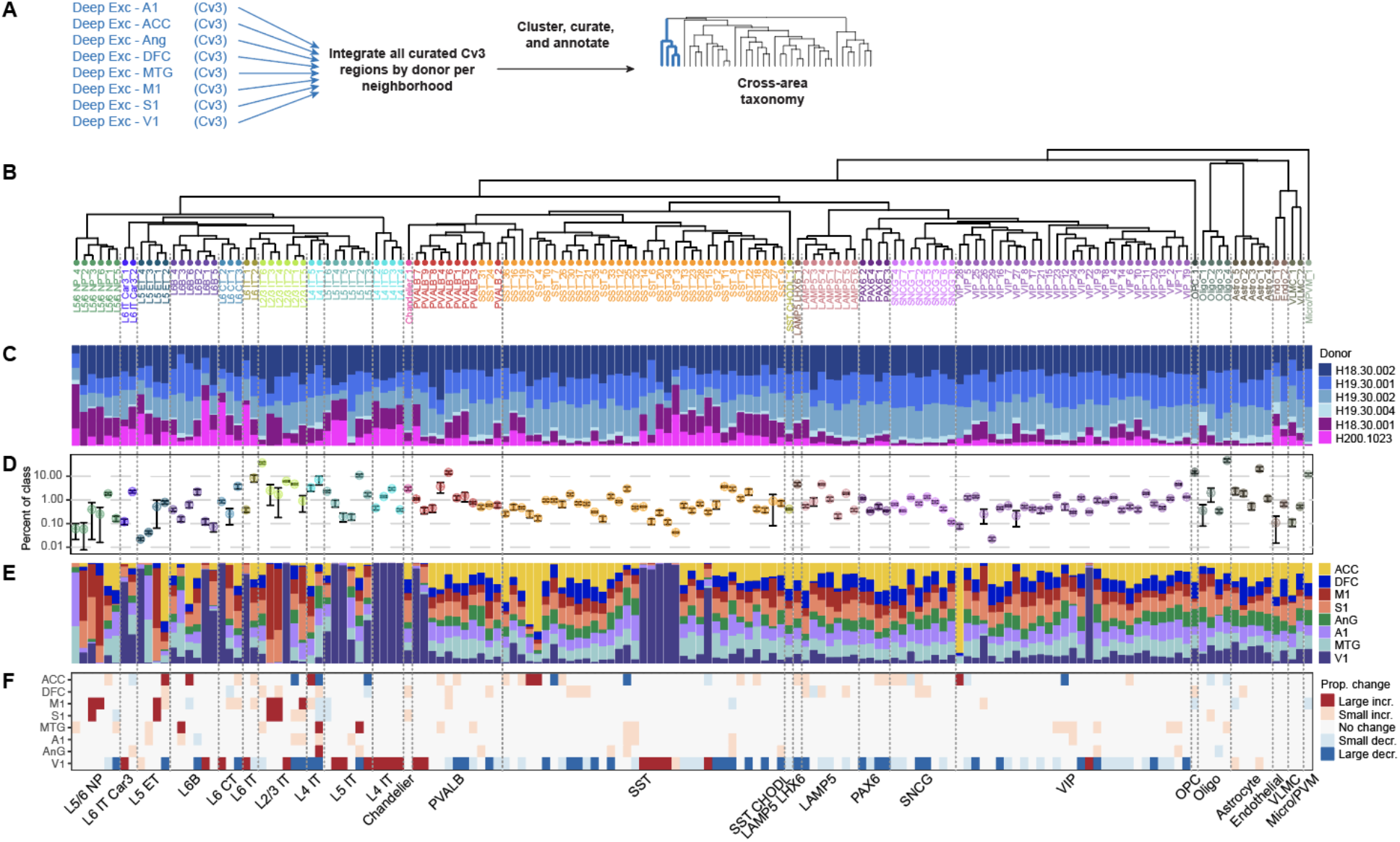
Cross-areal consensus taxonomy. **(A)** Schematic of data integration across donors used for each neighborhood (e.g. deep excitatory neurons) to generate the cross-area consensus taxonomy. **(B)** Consensus taxonomy of cell types across eight areas. **(C)** Proportion of nuclei in each consensus type dissected from each donor. **(D)** Consensus type proportion including nuclei from all areas as a fraction of cell class. Error bars indicate mean and standard deviation across donors. **(E)** The relative number of nuclei dissected from areas that contribute to each consensus cell type. **(F)** Significant changes in consensus type proportions across areas based on compositional analyses of neurons and non-neuronal cells using scCODA. Larger magnitude changes are indicated by darker colors. See **Table S10** for proportion effect sizes.

To determine significant changes in the relative abundances of cell types while accounting for the compositional nature of the data, we applied a Bayesian model (scCODA) to the snRNA-seq data. Grouping nuclei by consensus types and iteratively testing for consistent differences using each type as the “unchanged” reference population, all subclasses included consensus types with both increased and decreased proportions (**Table S10**), with the exception of PAX6 inhibitory types that had uniformly decreased abundances in V1. V1 had the most consensus types with consistent abundance changes (92 of 153, 60%), including two types with the largest changes (L4 IT_5 and L2/3 IT_2). While excitatory neurons were the most specialized in V1, several SST, PVALB, and VIP consensus types were also specific to V1. Specialized types were also found in other areas, including L2/3 IT and L5/6 NP excitatory types (L2/3 IT_3, L2/3 IT_4, L5/6 NP_3 and L5/6 NP_6) in M1 and S1, SST types (SST_4 and SST_10) in ACC, and distinct L5 ET types across the rostrocaudal axis. These dramatic changes, along with more subtle abundance changes (median 17 consensus types were affected in each area), likely contribute to the specialized functional role of each area.

### V1 specializations

The striking distinctiveness of V1 was reflected in the transcriptomic uniqueness of specific cell types. Considering cell types with >60% membership in V1 compared to other areas to be V1-specialized, there were specialized cell types in every excitatory subclass except L5/6 NP, with the greatest number of V1-specialized types in the L2/3 IT and L4 IT subclasses (**Fig. 6A** and **Table S11**). Surprisingly, given prior reports of common GABAergic neurons across the mouse neocortex (*15*, *19*), V1 had a number of specialized CGE- and MGE-derived types.

**Fig. 6.**
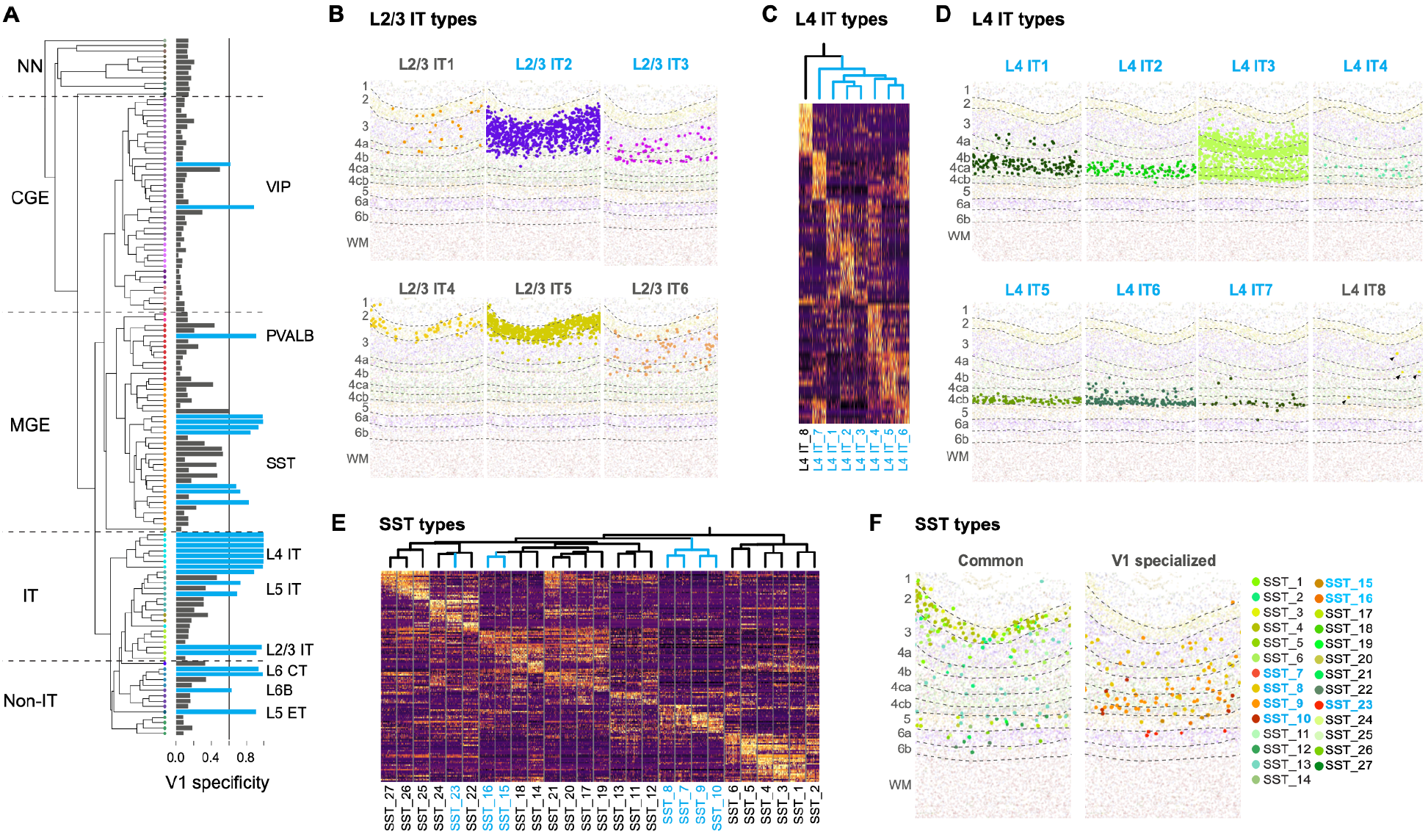
V1 cell type specialization. **(A)** Transcriptional uniqueness of cell types in the V1 taxonomy. Cell types with specificity >0.6 are considered V1-specialized and are highlighted in blue (see **Table S11**). **(B)** Laminar distributions of specialized (blue text) and common (grey) L2/3 IT types based on MERFISH *in situ* labeling experiments. **(C,E)** Scaled expression of marker genes of V1 specialized (blue labels) and common (black) L4 IT (C) and SST (E) types across areas. Dendrograms were pruned from the V1 taxonomy in panel A. **(D,F)** Laminar distributions of specialized and common L4 IT (D) and SST (F) types based on MERFISH experiments. L4 IT8 neurons are labeled by arrowheads.

MERFISH analysis of V1 provided the spatial organization of all cell types (**Fig. S11A,B**). The L2/3 IT types showed distinct markers (**Table S12**) and spatial distributions in terms of sublaminar organization and relative proportions (**Fig. 6B**). For example, L2/3 IT5 and L2/3 IT2 clearly delineated layer 2 and layer 3 from one another, respectively. Other L2/3 IT types were more sparsely distributed in layer 2 (L2/3 IT4), layer 3 (L2/3 IT3), or both layers 2 and 3 (L2/3 IT1 and 6). Interestingly, multiple L2/3 IT types were also found in layer 4A (e.g. L2/3 IT2) and even the superficial part of layer 4B (L2/3 IT3), and these types were V1-specialized. Conversely, the distributions of several L4 IT types were found in layers 4A and 4B and even into the deep part of layer 3 (e.g. L4 IT1 and 3, **Fig. 6D**). Thus, the specialized layers 4A and B contain not only L4 IT-type neurons, but also L2/3 IT-type neurons. This finding may help resolve ongoing questions about primate V1 layer 4A and 4B, which contains both stellate (L4 IT-like) and pyramidal corticocortical projection neurons (L2/3 IT-like) (*33*).

Layer 4 of primary visual cortex, or striate cortex, is highly distinctive even in unlabeled tissues, due to the band of myelinated thalamocortical axons entering layer 4 that form the stria of Gennari. This distinctiveness was also seen at the level of L4 IT neuron types, all but one of which were V1-specialized (**Fig. 6C, D**). As in layers 2 and 3, different L4 IT types showed different markers (**Table S12**) and sublaminar distributions, from dense pan-layer 4 (L4 IT3) to sublayer-specific distributions. Layers 4C*α* and 4C*β* receive selective inputs from magnocellular and parvocellular layers of the thalamic lateral geniculate nucleus, respectively. Corresponding to these functionally segregated visual inputs, selective localization of specific types was found in each sublayer. L4 IT5 was selectively localized in layer 4C*β*, while L4 IT2 was enriched in layer 4C*α* but extending into layer 4B. This latter localization is consistent with the fuzzy boundary between 4C*α* and 4B described in other human studies (*34*). Other sparser L4 IT types were scattered across layers. Together these illustrate the cellular specialization of the distinctive input layer of V1, and a complexity of putative thalamorecipient stellate neurons that offers many avenues for future exploration.

L6 CT neurons that send reciprocal projections to the LGN were also highly specialized in V1 (**Fig. 6A**), with two unique types that expressed many V1-enriched genes that suggest significant specialization of physiological and connectional properties (**Fig. S11C**). Gene set enrichment analysis showed highly significant enrichment for calcium signaling and axon guidance and axonal and synaptic compartments (**Fig. S11D**). For example, axon guidance molecules *CDH7, EPHA6*, and *SEMA6A* showed V1 enrichment, the latter previously shown to have V1-selective developmental expression presumably involved in establishing connections to the lateral geniculate nucleus (*35*). Various ion channels (e.g. *KCNT2* and *SCN1B*) and synaptic genes (e.g. *SYT6* (*36*)), as well as calcium and calmodulin signaling associated genes (e.g. *PCP4* (*37*), *NPY2R* (*36*)) were similarly enriched with several showing conserved V1 enrichment in monkey (*36*), as well as myelin basic protein (*MBP*), normally described in oligodendrocytes but known to function in certain neurons as part of a Golli-MBP complex, including in developing human visual cortex where it is hypothesized to play a role in developmental plasticity (*38*).

In addition to specialized excitatory neurons, V1 also contained specialized GABAergic interneuron types. The bulk of these were SST types (**Fig. 6E, F** and **Table S12**), and also one PVALB and two VIP types (**Fig. S11A**). Interestingly, the SST types common across V1 and other areas were concentrated in layer 2 with sparser representation in other layers. In contrast, the V1-specialized types were concentrated in layer 4, in close proximity to the V1-specialized L4 IT types, suggestive of a relationship between these specialized excitatory and inhibitory types as discussed below.

### L5 ET neuron diversity

Neocortical extratelencephalic (subcerebral) projecting axons originate from excitatory neurons in layer 5 that have distinctive electrophysiological properties and gene expression patterns (*12*, *39*). The hallmark morphologies of these cells have historically been key defining features of the cytoarchitecture of cortical areas, with M1 characterized by the presence of gigantopyramidal Betz cells and ACC (along with the frontoinsular cortex) characterized by spindle-shaped von Economo neurons (VENs), for example (**Fig. 7A**). Capturing these sparse neuron populations required additional sampling with 10x Cv3 on dissected layer 5 samples. As noted above, L5 ET neurons were most abundant in ACC and their abundance generally decreased along the rostrocaudal axis (**Fig. 2D**). V1 had the lowest proportion of L5 ET neurons (^~^0.1% of excitatory neurons), consistent with data from macaque monkeys demonstrating projections to subcortical targets such as the superior colliculus from large, very sparse neurons localized to deep layers in V1 (e.g., Meynert cells, **Fig. S12A**) (*40–42*).

**Fig. 7.**
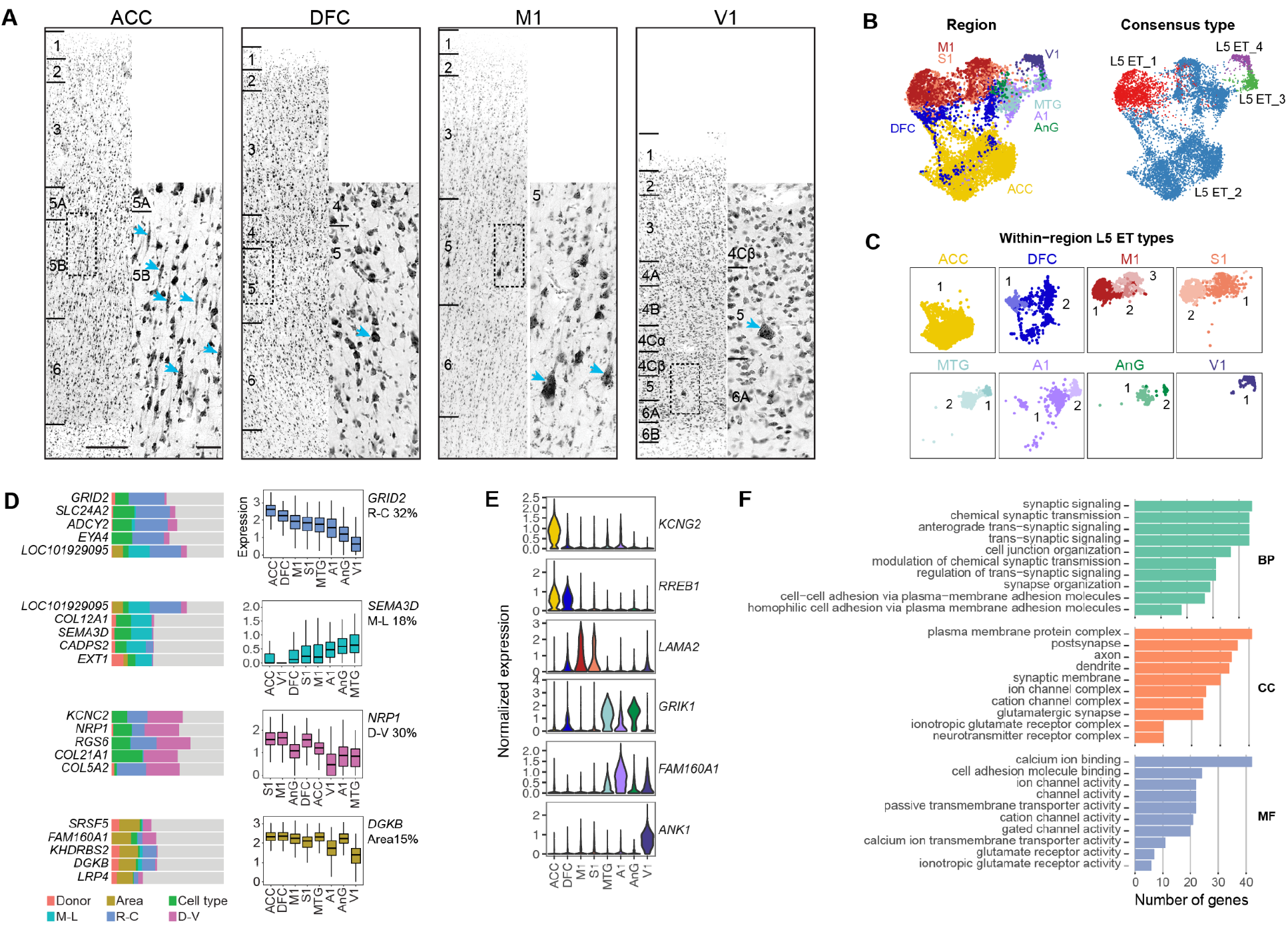
L5 ET-projecting neuronal diversity. **(A)** L5 ET morphology varies across areas based on fluorescent Nissl stains. **(B)** UMAPs of L5 ET neurons labeled by area and cross-area consensus type. **(C)** Within-area L5 ET subtypes for each area shown in the same UMAP space as panel B. **(D)** Bar plots summarizing the expression variance explained by human donor, L5 ET subtype, and four types of variation across areas: rostrocaudal (R-C), mediolateral (M-L), and dorsoventral (D-V) gradients and more complex patterns or in a single area (Area). For the four types of areal variation, the distribution of expression across areas is shown for one of the top five genes. **(E)** Examples of genes with L5 ET neuron expression restricted to one or a few areas. **(F)** Number of genes in the top 10 significantly enriched terms from Gene ontology (GO) analyses (biological process, BP; cellular component, CC; molecular function, MF) of L5 ET areal markers (**Table S13**). Scale bars on A = 250μm and 50μm (inset).

We identified 4 consensus L5 ET types (**Fig. 5; Fig. 7B**), several of which were dominated by nuclei derived from one or two cortical areas in close proximity to each other. For example, M1 and S1 predominantly contributed to L5 ET_1, whereas L5 ET_3 was largely composed of nuclei from nearby areas MTG and A1 (and to a lesser extent AnG), again suggesting similarity in gene expression signatures based on the topographic position of cortical areas. V1 specialization was also apparent in L5 ET consensus types, with only a single type, L5 ET_4, consisting of nuclei almost exclusively isolated from V1 (**Fig. 7B**). L5 ET neurons could be further divided into at least two transcriptomically distinct subtypes in most regions (**Fig. 7C**), and M1 had 3 distinct subtypes as reported in our previous study (*12*) where we determined that at least 2 L5 ET M1 subtypes included Betz cells. Interestingly, despite having the highest proportion of L5 ET neurons of all cortical areas, only one subtype was identified in ACC, suggesting that VENs in ACC likely do not represent a distinct transcriptomic cluster but rather are transcriptionally similar to other L5 ET morphological types, consistent with our findings in frontoinsula (*43*).

L5 ET neurons have more genes with variable expression across areas than any other cell type (**Fig. S10F**). Up to 32% of variation in gene expression across areas was explained by gradients along mediolateral, dorsoventral, and rostrocaudal axes (**Fig. 7D**). Top gradient genes included a glutamate receptor subunit (*GRID2*), a semaphorin (*SEMA3D*), and a neuropilin (*NRP1*) that are involved in trans-synaptic signaling and connectivity (**Fig. 7D**). Some gene expression varied across areas but not as a gradient, such as *DGKB*, which was selectively down-regulated in primary sensory areas (A1, S1, V1) and has been shown to regulate spine formation of medium spiny neurons in the striatum (*44*). L5 ET neurons also expressed distinct areal markers (**Fig. 7E** and **Table S13**), including the voltage-gated potassium channel *KCNG2* in ACC, glutamate receptor subunit *GRIK1* in MTG and AnG, and *ANK1* in V1, a gene that encodes for the scaffolding protein Ankyrin 1 and was shown to be enriched in mouse cerebellar neurons (*45*). Gene ontology (GO) enrichment analysis of L5 ET areal markers identified significantly enriched pathways associated with synaptic signaling, connectivity, and intrinsic neuronal firing properties (**Fig. 7F**), consistent with known areal variation in firing properties.

### Glial specialization

Non-neuronal cells comprised at least 40-65% of cortical cells across areas based on FACS analysis of dissociated nuclei labeled with the neuronal marker NeuN and gated based on NeuN fluorescence intensity (**Fig. S12A**). However, these proportions are an underestimate of the total non-neuronal population because vascular cells, including endothelial cells and VLMCs, are difficult to dissociate (*46*) and are undersampled in the snRNA-seq dataset based on *in situ* labeling with MERFISH (**Fig. S12B**) (*17*). M1 and S1 had a higher proportion of non-neuronal (NeuN-negative) cells than other areas by FACS, and snRNA-seq data showed this was driven by an expansion of oligodendrocytes relative to OPCs, astrocytes, and microglia (**Fig. S12B,C,D**), consistent with previous neuroimaging studies showing that these areas are the most heavily myelinated of all cortical areas (*47*, *48*) and with the presence of heavily myelinated axons of deep excitatory neurons in these areas (*49*). By contrast, areas previously described to be among the most lightly myelinated in cortex (ACC and DFC) (*47*) had the lowest proportion of oligodendrocytes (**Fig. S12B,C**).

Non-neuronal cells were grouped into major subclasses based on conserved marker expression across cortical areas (**Fig. S12F**), and many subclasses could be further divided into transcriptomically distinct subtypes. Astrocytes could be subdivided into previously described protoplasmic, interlaminar (ILM), and fibrous types, which also had robust markers across areas (**Fig. S12G**). Consistent with previous reports of shared non-neuronal types across cortical areas (*15*), there was little areal expression signature for most non-neuronal cell types (**Fig. 8A**, **Fig. S12E**). By contrast, areal variation in protoplasmic, but not ILM or fibrous astrocytes, was apparent and this was consistent with previous reports describing brain-wide astrocyte heterogeneity (*20*, *50*) and variation in astrocytes across cortical and hippocampal areas in mouse (*51*). Interestingly, protoplasmic astrocytes from ACC grouped together on the UMAP and expressed distinct areal markers (*NRP2, NR4A3*, and *LGR6*) (**Fig. 8B**).

**Fig. 8.**
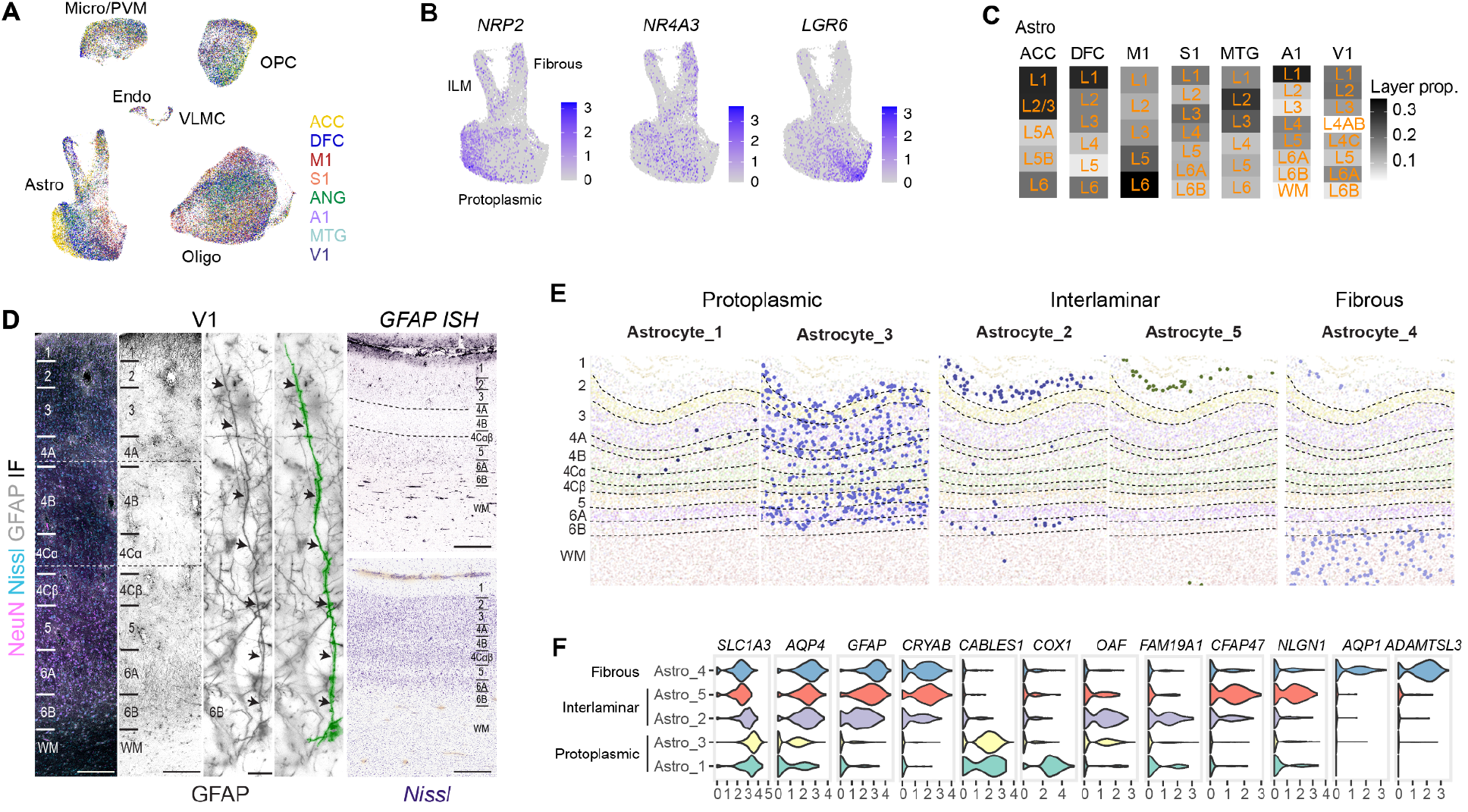
Areal specialization of astrocytes. **(A)** UMAP of non-neuronal cells labeled by cortical area. **(B)** UMAPs of astrocyte expression for genes with areal enrichment. **(C)** Laminar distributions of astrocytes vary across areas. **(D)** GFAP immunofluorescence (IF) and *in situ* hybridization (ISH) illustrates variable laminar distributions and morphologies of astrocytes in V1. Single channel IF images were inverted to increase visibility of GFAP IF. Scale bars: IF columns (100μm), GFAP tracing images (15 μm), ISH (200 μm). **(E)** Laminar distributions of astrocyte subtypes in V1 based on MERFISH *in situ* labeling experiments. **(F)** Pan-astrocyte and subtype marker expression.

Laminar distributions varied across areas for all glial subclasses (**Fig. 8C, S12H**). In particular, there was a striking depletion of astrocytes in L4A and L4B of V1 but not in L4 of other sensory or granular cortical areas (**Fig. 8C**). To validate this finding, we examined *in situ* expression of the astrocyte marker GFAP in V1 and another granular area, DFC. We confirmed that GFAP protein and gene expression was reduced in L4B of V1 (**Fig. 8D**), and only protein expression was reduced in L4 of DFC (**Fig. S12I**) based on immunofluorescence (IF) and *in situ* hybridization (ISH) labeling of adult human tissue. In V1, a band of dense GFAP labeling was apparent in layers 6A and 6B, which tapered off in the underlying white matter. Further examination of GFAP IF in V1 revealed a population of astrocytes that extended long GFAP-positive processes away from the white matter and into L5, similar to descriptions of varicose projection astrocytes (VPA) that are unique to humans and great apes and not found in the cortex of other anthropoid primates (*52*, *53*) (**Fig. 8D**, **S12I**). Deep layer astrocytes in DFC did not extend long processes and had morphology typical of protoplasmic and fibrous astrocytes (**Fig. S12I**).

To investigate the transcriptomic identities of astrocytes in V1 further, we used MERFISH to map the spatial distributions of the five astrocyte subtypes identified in V1 by snRNA-seq (**Fig. S8**). Based on laminar distributions (**Fig. 8E**) and marker gene expression (**Fig. 8F**), there were two subtypes of protoplasmic (Astro_1 and Astro_3) and ILM (Astro_2 and Astro_5) astrocytes and one fibrous subtype (Astro_4). Contrary to prior descriptions of protoplasmic astrocytes as relatively homogenous cells, protoplasmic subtypes in V1 displayed distinct laminar patterns with Astro_1 localized predominantly to the sublayers of layer 4 and Astro_3 spread across layers 2-6 but strikingly absent in layers 1, 6B and white matter. Astro_1 markers were related to energy metabolism, including mitochondrial genes *COX1* (**Fig. 8F**), *COX2*, and *COX3*, and Astro_1 cells may be highly active protoplasmic astrocytes (i.e. an Astro_3 cell state) rather than a developmentally distinct type. Astro_5 cells were almost entirely restricted to the layer 1-pial border, while Astro_2 cells were enriched in the deeper part of layer 1, and these subtypes likely represent pial and subpial ILMs (*54*), respectively. Interestingly, the putative subpial ILM type (Astro_2) included a small number of cells localized to deep L6. Since ILMs and VPAs have previously been shown to express shared marker genes (e.g. *AQP4* and *CRYAB*, **Fig. 8F**) and have similar morphologies (*53*, *55*), these deep layer Astro_2 cells may represent a type of VPA. However, further work will be needed to fully characterize the diversity of astrocyte morphologies across cortex and their relationships to transcriptomic astrocyte types.

## Discussion

The cellular complexity of the neocortex has challenged generations of neuroscientists aiming to understand the structural basis of cognitive function. Single cell genomics has rapidly matured to provide the scale and information content to quantitatively define cellular diversity and map the spatial tissue organization of these cell types in a remarkably comprehensive fashion. The BRAIN Initiative Cell Census Network established a paradigm for mapping cortical cellular diversity, developed methods to work across species including human, and established the concordance of a transcriptomic cellular classification with other cellular properties in a way that integrates a vast prior literature while identifying a much greater cellular diversity than appreciated with prior studies (*12*, *16*). Among the lessons learned from those studies is that most cortical cell types are rare, but can be aggregated hierarchically in subclasses that reflect major divisions with laminar, developmental and projection target consistency. Characterizing this full diversity requires deep cellular coverage (80-100k nuclei per area), and very rare cell types like L5 ET may require additional sampling or selection to adequately represent. Furthermore, co-expression patterns of cortical cells are sufficiently conserved across mammals to align and transfer cell type labels from a well annotated cortical area in mouse to homologous cell types across species or to other cortical areas.

Here we used these principles to perform cross-areal analysis across a series of human cortical areas, building on our highly annotated M1 taxonomy to understand similarities, differences and organizational principles. Since the neocortex has a common organization and also graded changes and areal specializations, we applied two complementary analysis strategies to define cell types. The first strategy was to analyze each area independently, transferring labels from the most highly annotated area (M1 in this case) to other areas. The advantage of this strategy is that it provides the highest resolution clustering in each area and clearly identifies a common subclass-level organization. The other strategy was to analyze data from all areas jointly, identifying a set of consensus clusters that may be present in multiple areas, but that also captures cell types that are unique to a single area. This latter approach more clearly illustrates common cell types across areas (especially neighboring areas) and highly specialized types. Each of these approaches illustrate different aspects of cortical organization, which has aspects of areal uniqueness and also graded effects that clustering algorithms may attempt to split when those differences along the gradient become large. Similar joint analysis strategies have been used recently on the entire mouse cortex with similar results (*19*).

A key finding is that all 24 subclasses identified in M1 are found in all other cortical areas, substantiating the idea that there is a common organization across the entire neocortex at that level of granularity. This was true for L4 IT-like neurons, which were found even in agranular ACC and M1 (*12*) despite the lack of a histologically obvious layer 4. Each cortical area analyzed could be defined as a unique proportional makeup of cell subclasses. The majority of proportional variation was due to variation in excitatory neuron subclasses, which could be dramatic (10 to 50-fold), in particular in L4 IT and L5 ET neurons. At a finer level of analysis, there was a substantial areal variation such that distant areas had distinct cellular expression and some cell types clustered separately. Thus, both a canonical and a non-canonical architecture were apparent, depending on the level of cellular detail analyzed.

Topographic variation as a function of rostrocaudal position was an obvious organizational feature. Prior microarray-based analysis of human (*30*) and macaque (*36*) cortex has shown molecular similarity varies as a function of distance on the cortical sheet, strongly reflecting topographic organization that likely mirrors early developmental gradients of transcription factors and morphogens (*56*, *57*). Here we see similar variation by rostrocaudal position and similarity as a function of distance, but predominantly in select cell types. As in mouse cortex (*15*), most areal variation was in the excitatory neuron populations and not the inhibitory neuron populations (except V1 as discussed below). These results are consistent with the fact that most inhibitory neurons migrate in from the ganglionic eminences and are complex but relatively homogeneous across the cortex, whereas excitatory neurons are generated from progenitor cells with strong developmental gradients that are maintained in postmitotic neurons in the adult cortex. Interestingly, rostrocaudal variation was seen not just in gene expression characteristics but also in excitatory neuron proportions. L4 IT neuron proportions appeared to increase from rostral to caudal, while L5 ET neurons proportions were inversely correlated. These observations suggest that there are developmental processes based on early gradients that sculpt cortical cellular makeup in many ways. Similar rostrocaudal variation in cellular morphology in primate species supports this idea as well (*58*).

The most dramatic observation was the distinctiveness of V1, mirroring the specialized cytoarchitecture of V1 in human, primates and other binocular mammals. Unlike in mouse cortex where VISp is not highly different from other cortical areas (*19*), V1 in human was an outlier in terms of molecular distinctiveness, cellular proportions and cell type makeup. V1 was more molecularly distinctive than expected by topographic position, as shown previously with bulk microarray analysis (*28*), with most excitatory neuron types appearing as separate islands on UMAP representations. L4 IT proportions were dramatically expanded, as expected given the increased size of the thalamorecipient layer 4. In contrast, L5 ET neurons in V1, presumably containing Meynert cells, were by far the most sparse among cortical areas. The corticothalamic L6 CT neurons were extremely distinctive from those of other areas, including the other primary sensory areas S1 and A1. At the more granular cell type level, V1 had many unique cell types. This included 7 layer 4 types in Layer 4A, 4B and 4C*α/β* that may reflect different thalamic input specificity known to vary by layer 4 sublayer. However, every excitatory neuron subclass except L5 NP had a V1-specific type; furthermore, there were V1-specific SST and PVALB types. Interestingly, the V1-specific SST types were found in layer 4 as well, suggesting either converging evolutionary or developmental sculpting of the excitatory and inhibitory neuron types.

Finally, there has been long-standing debate about the cellular makeup of V1 layers 4A and 4B, which have alternatively been called 3BP and 3C in the Hassler nomenclature (*33*, *59*). Cells in 4A and 4B have features of both layer 3 long-range projection neurons and also thalamorecipient local circuit spiny stellate cells normally found in layer 4 (and spiny stellate projection neurons not seen in other areas). Cytochrome oxidase labeling in human V1 that typically labels thalamocortical afferent termination is also inconclusive, failing to show a clear 4A or a clear 4C*α*/4B border (*60*). Our results provide an explanation for this confusion: Layers 4A and 4B contain both L4 IT-like neuron types and also L2/3 IT-like neuron types, in particular with a V1-specific L2/3 IT neuron type that extends into layer 4. Additional cellular diversity that does not strictly obey laminar boundaries complicates this organization, similar to previous work showing lack of strictly laminar cell organization in human MTG (*11*). These data now provide both an explanation and a map to guide characterization of the properties of the various V1 cell types using Patch-seq methods in monkey cortex, which is likely to have highly conserved cellular makeup to human.

The balance of excitation and inhibition is thought to be critical to proper balance of neuronal circuitry, with disruption of E:I balance as a potential mechanism underlying epilepsies, neurodevelopmental and neuropsychiatric diseases(*61*). Reported ratios based on immunohistochemistry for GABA have been widely ranging from ^~^4:1 in human frontal cortex (*62*) to 4:1 in monkey V1 (*63*, *64*). Transcriptomic analysis offers a more reliable method than antibody labeling to define and quantify cell proportions, and recent reports had shown a significant species variation in the cortical E:I ratio of excitatory to inhibitory neurons (*12*). Whereas this ratio is about 5:1 in mouse cortex, the ratio in human MTG and M1 is closer to 2:1. This finding was confirmed by MERFISH here and in (*17*), and also by analysis of EM volumes of mouse and human layer 2 (*65*), although the latter finding that there may be homeostatic processes that achieve similar synaptic E:I balance despite differences in cell numbers. We find that the human E:I ratio of 2:1 is consistent across all areas except V1, where the ratio is 5:1, similar to mouse. This is likely due largely to the increase in L4 IT neurons in V1 as a whole. However, we observed that this ratio varies substantially by layer as well, with the laminar variation most dramatic in V1 where the E:I ratio in layer 6 is closer to 10:1. Whether this variation can be compensated by homeostatic processes remains to be studied, but these results indicate that the E/I ratio can vary quite dramatically in human cortex, with both laminar variation and V1-specific areal variation.

The current results illustrate the power of single cell genomics to provide a comprehensive cellular map of the cortex that can be thought of as a new form of quantitative cytoarchitectonics based on the genes that give the cell types their properties. These analyses provide a new map for the field to understand organizational principles and place a cellular lens on thinking about cortical functional variation as variation in the proportions and properties of the component cell types that define the input-output properties of those areas. Recent studies have shown that morphological and anatomical characteristics are correlated with transcriptomic identity and gradient properties (*16*, *24*, *66*), indicating that the transcriptomic maps are also highly predictive for cell phenotype variation. Challenges for the future will be to map the entire human neocortex, understand graded features versus discrete boundaries, and directly measure the relationship between transcriptomically defined cell types, cellular phenotypes and functional architecture.

## Methods

### Post-mortem tissue donors

Males and females 18 – 68 years of age with no known history of neuropsychiatric or neurological conditions (‘control’ cases) were considered for inclusion in this study. De-identified postmortem human brain tissue was collected after obtaining permission from the decedent’s legal next-of-kin. Tissue collection was performed in accordance with the provisions of the United States Uniform Anatomical Gift Act of 2006 described in the California Health and Safety Code section 7150 (effective 1/1/2008) and other applicable state and federal laws and regulations. The Western Institutional Review Board (WIRB) reviewed the use of de-identified postmortem brain tissue for research purposes and determined that, in accordance with federal regulation 45 CFR 46 and associated guidance, the use deidentified specimens from deceased individuals did not constitute human subjects research requiring IRB review. Routine serological screening for infectious disease (HIV, Hepatitis B, and Hepatitis C) was conducted using donor blood samples and donors negative for all three infectious diseases were considered for inclusion in the study. Tissue RNA quality was assessed using samples of total RNA derived from the frontal and occipital poles of each donor brain which were processed on an Agilent 2100 Bioanalyzer using the RNA 6000 Nano kit to generate RNA Integrity Number (RIN) scores for each sample. Specimens with average RIN values ≥7.0 were considered for inclusion in the study.

### Processing of whole brain postmortem specimens

Whole postmortem brain specimens were transported to the Allen Institute on ice and processed as previously described (https://dx.doi.org/10.17504/protocols.io.bf4ajqse). Briefly, brain specimens were bisected through the midline and individual hemispheres were embedded in Cavex Impressional Alginate for slabbing. Coronal brain slabs were cut at 1 cm intervals and individual slabs were frozen in a slurry of dry ice and isopentane. Frozen slabs were vacuum sealed and stored at −80°C until the time of use.

### Tissue microdissection, nucleus isolation and capture

Cortical areas of interest were identified on tissue slab photographs taken at the time of autopsy and were removed from frozen slabs held on a −20°C custom cold table during dissection. Dissections of DFC were targeted to the superior frontal gyrus corresponding to the lateral and medial subdivisions of Brodmann Area (A) 9. Dissections of ACC corresponded to A24 in the rostral (anterior) cingulate gyrus. A1 was localized in the transverse temporal gyrus (Heschl’s gyrus) corresponding approximately to A41. For M1 and S1 dissections, putative hand and trunk-lower limb sub-regions of each cortical area were identified on tissue photographs, removed from slabs of interest, and subdivided into smaller blocks. One block from each donor was processed for cryosectioning and fluorescent Nissl staining (Neurotrace 500/525, ThermoFisher Scientific) and stained sections were screened for histological hallmarks of each cortical area (e.g., the presence of Betz cells in L5 of M1) to verify that dissected regions were appropriately localized to either M1 or S1. MTG sampling targeted predominantly the caudal subdivision of A21 and the area of AnG sampled corresponded approximately to A39 located lateral to the intraparietal sulcus. V1 was grossly identifiable on tissue slab photographs by the presence of the Stria of Gennari.

For SMART-seqv4 (SSv4) processing, tissue blocks were placed in ice-cold 1X PBS supplemented with 10mM DL-Dithiothreitol (DTT, Sigma Aldrich) and mounted on a vibratome (Leica) for sectioning at 500μm in the coronal plane. Sections were placed in fluorescent Nissl staining solution (Neurotrace 500/525, ThermoFisher Scientific) prepared in 1X PBS with 10mM DTT and 0.5% RNasin Plus RNase inhibitor (Promega) and stained for 5 min on ice. After staining, sections were visualized on a fluorescence dissecting microscope (Leica) and cortical layers were individually microdissected using a needle blade micro-knife (Fine Science Tools) as previously described (https://dx.doi.org/10.17504/protocols.io.bq6ymzfw). Nuclear suspensions were prepared from microdissected tissue pieces as described (https://dx.doi.org/10.17504/protocols.io.ewov149p7vr2/v2). For 10xv3 (Cv3) processing, tissue blocks encompassing all cortical layers were placed directly into a Dounce homogenizer after removal from the −80°C freezer and processed as described (https://dx.doi.org/10.17504/protocols.io.bq64mzgw). For L5 specific dissections, tissue blocks were sectioned using a vibratome as described above and L5 was specifically dissected from Nissl stained vibratome sections using a micro-knife. Dissected L5 tissue pieces were pooled across multiple sections per cortical area and were processed for nuclear isolation as described above.

All samples were immunostained for fluorescence activated cell sorting (FACS) with mouse anti-NeuN conjugated to PE (EMD Millipore, FCMAB317PE) at a dilution of 1:500 with incubation for 30 min at 4°C. Control samples were incubated with mouse IgG1,k-PE Isotype control (BD Pharmingen). A subset of SSv4 samples was immunostained with rabbit anti-SATB2 conjugated to Alexa Fluor 647 (Abcam, ab196536) at a dilution of 1:500 to discriminate excitatory (SATB2+/NeuN+) and inhibitory (SATB2−/NeuN+) nuclei. After immunostaining, samples were centrifuged to concentrate nuclei and were resuspended in 1X PBS, 1% BSA, and 0.5% RNasin Plus for FACS. DAPI (4’, 6-diamidino-2-phenylindole, ThermoFisher Scientific) was applied to samples at a concentration of 0.1μg/ml. Single nucleus sorting was carried out on either a BD FACSAria II SORP or BD FACSAria Fusion instrument (BD Biosciences) using a 130μm nozzle. A standard gating strategy was applied to all samples as previously described (Hodge et al., 2019). Briefly, nuclei were gated on their size and scatter properties and then on DAPI signal. Doublet discrimination gates were applied to exclude multiplets. Lastly, samples were gated on NeuN signal (PE) and SATB2 (Alexa Fluor 647) signal where applicable. For Cv3 experiments, NeuN+ and NeuN-nuclei were sorted into separate FACS tubes and combined at defined ratios (90% NeuN+, 10% NeuN-), except for L5 dissected samples where only neuronal (NeuN+) nuclei were captured. Samples were then centrifuged and resuspended in 1XPBS, 1% BSA, 0.5% RNasin Plus, and 5-10% DMSO and frozen at −80°C until the time of chip loading. Samples were processed according to the following protocol for chip loading (https://dx.doi.org/10.17504/protocols.io.774hrqw). For SSv4, single nuclei were sorted into 8-well strip tubes containing 11.5μl of SMART-seq v4 collection buffer (Takara) supplemented with ERCC MIX1 spike-in synthetic RNAs at a final dilution of 1×10-8 (Ambion). Strip tubes containing sorted nuclei were briefly centrifuged and stored at −80°C until the time of further processing.

### SMART-seqv4 RNA-sequencing

We used the SMART-Seq v4 Ultra Low Input RNA Kit for Sequencing (Takara #634894) per the manufacturer’s instructions for reverse transcription of RNA and subsequent cDNA amplification as described (https://dx.doi.org/10.17504/protocols.io.8epv517xdl1b/v2). Standard controls were processed alongside each batch of experimental samples. Control strips included: 2 wells without cells, 2 wells without cells or ERCCs (i.e. no template controls), and either 4 wells of 10 pg of Human Universal Reference Total RNA (Takara 636538) or 2 wells of 10 pg of Human Universal Reference and 2 wells of 10 pg Control RNA provided in the Clontech kit. cDNA was amplified with 21 PCR cycles after the reverse transcription step. cDNA libraries were examined on either an Agilent Bioanalyzer 2100 using High Sensitivity DNA chips or an Advanced Analytics Fragment Analyzer (96) using the High Sensitivity NGS Fragment Analysis Kit (1bp-6000bp). Purified cDNA was stored in 96-well plates at −20°C until library preparation.

The NexteraXT DNA Library Preparation (Illumina FC-131-1096) kit with NexteraXT Index Kit V2 Sets A-D (FC-131-2001, 2002, 2003, or 2004) was used for sequencing library preparation as described (*11*). NexteraXT DNA Library prep was done at either 0.5x volume manually or 0.4x volume on the Mantis instrument (Formulatrix, https://dx.doi.org/10.17504/protocols.io.brdjm24n). Samples were quantitated using PicoGreen on a Molecular Bynamics M2 SpectraMax instrument. Sequencing libraries were assessed using either an Agilent Bioanalyzer 2100 with High Sensitivity DNA chips or an Advanced Analytics Fragment Analyzer with the High Sensitivity NGS Fragment Analysis Kit for sizing. Molarity was calculated for each sample using average size as reported by Bioanalyzer or Fragment Analyzer and pg/μl concentration as determined by PicoGreen. Samples were normalized to 2-10 nM with Nuclease-free Water (Ambion). Libraries were multiplexed at 96 samples/lane and sequenced on an Illumina HiSeq 2500 instrument using Illumina High Output V4 chemistry.

### SMART-seqv4 RNA-seq gene expression quantification

Raw read (fastq) files were aligned to the GRCh38 human genome sequence (Genome Reference Consortium, 2011) with the RefSeq transcriptome version GRCh38.p2 (current as of 4/13/2015) and updated by removing duplicate Entrez gene entries from the gtf reference file for STAR processing. For alignment, Illumina sequencing adapters were clipped from the reads using the fastqMCF program (*67*). After clipping, the paired-end reads were mapped using Spliced Transcripts Alignment to a Reference (STAR)(*68*) using default settings. Reads that did not map to the genome were then aligned to synthetic constructs (i.e. ERCC) sequences and the E.coli genome (version ASM584v2). The final results files included quantification of the mapped reads (raw exon and intron counts for the transcriptome-mapped reads), and percentages of reads mapped to the RefSeq transcriptome, to ERCC spike-in controls, and to E.coli. Quantification was performed using summerizeOverlaps from the R package GenomicAlignments (*69*).

Expression levels were calculated as counts per million (CPM) of exonic plus intronic reads, and log2(CPM + 1) transformed values were used for a subset of analyses as described below. Gene detection was calculated as the number of genes expressed in each sample with CPM > 0. CPM values reflected absolute transcript number and gene length, i.e. short and abundant transcripts may have the same apparent expression level as long but rarer transcripts. Intron retention varied across genes so no reliable estimates of effective gene lengths were available for expression normalization. Instead, absolute expression levels were estimated as fragments per kilobase per million (FPKM) using only exonic reads so that annotated transcript lengths could be used.

### 10x Chromium RNA-sequencing and expression quantification

Samples were processed using the 10x Chromium Single-Cell 3’ Reagent Kit v3 following the manufacturer’s protocol as described (https://dx.doi.org/10.17504/protocols.io.bq7cmziw). Gene expression was quantified using the default 10x Cell Ranger v3 (Cell Ranger, RRID:SCR_017344) pipeline. The human reference genome used included the modified genome annotation described above for SMART-seq v4 quantification. Introns were annotated as “mRNA” and intronic reads were included in expression quantification.

### RNA-sequencing processing and clustering

#### Cell type label transfer

Human M1 reference taxonomy subclass labels (*12*) were transferred to nuclei in the current MTG dataset using Seurat’s label transfer (3000 high variance genes using the ‘vst’ method then filtered through exclusion list). This was carried out for each RNA-seq modality dataset; for example, human-Cv3 and human-SSv4 were labeled independently. Each dataset was subdivided into 5 neighborhoods – IT and Non-IT excitatory neurons, CGE- and MGE-derived interneurons, and nonneuronal cells – based on marker genes and transferred subclass labels from published studies of human and mouse cortical cell types and cluster grouping relationships in a reduced dimensional gene expression space.

#### Filtering low-quality nuclei

SSv4 nuclei were included for analysis if they passed all QC criteria:

> 30% cDNA longer than 400 base pairs
> 500,000 reads aligned to exonic or intronic sequence
> 40% of total reads aligned
> 50% unique reads
> 0.7 TA nucleotide ratio

QC was then performed at the neighborhood level. Neighborhoods were integrated together across all areas and modality; for example, deep excitatory neurons from human-Cv3, human-Cv3-Layer5 and human-SSv4 datasets were integrated using Seurat integration functions with 2000 high variance genes. Integrated neighborhoods were Louvain clustered into over 100 meta cells, and Low-quality meta cells were removed from the dataset based on relatively low UMI or gene counts (included glia and neurons with greater than 500 and 1000 genes detected, respectively), predicted doublets (include nuclei with doublet scores under 0.3), and/or subclass label prediction metrics within the neighborhood (ie excitatory labeled nuclei that clustered with majority inhibitory or non-neuronal nuclei).

#### RNA-seq clustering

Nuclei were normalized using SCTransform (*70*), and neighborhoods were integrated together within an area and across individuals and modalities by identifying mutual nearest neighbor anchors and applying canonical correlation analysis as implemented in Seurat (*71*). For example, deep excitatory neurons from human-Cv3 were split by individuals and integrated with the human-SSv4 deep excitatory neurons. Integrated neighborhoods were Louvain clustered into over 100 meta cells. Meta cells were then merged with their nearest neighboring meta cell until merging criteria were sufficed, a split and merge approach that has been previously described (*12*). The remaining clusters underwent further QC to exclude Low-quality and outlier populations. These exclusion criteria were based on irregular groupings of metadata features that resided within a cluster.

### Defining cross-area consensus cell types

For each neighborhood, Cv3 nuclei were integrated together across individuals. The integrated latent space was Louvain clustered into over 100 meta cells. Meta cells were then merged with their nearest neighboring meta cell until merging criteria were sufficed, a split and merge approach that has been previously described (*12*) and was also used to define the within-area cluster identities. The process was repeated for each neighborhood, with an example diagram of the workflow shown in Figure 5A.

### Cell type taxonomy generation

For each area, a taxonomy was built using the final set of clusters and was annotated using subclass mapping scores, dendrogram relationships, marker gene expression, and inferred laminar distributions. Within-area taxonomy dendrograms were generated using build_dend function from scrattch_hicat R package. A matrix of cluster median log2(cpm + 1) expression across the 3000 High-variance genes for Cv3 nuclei from a given area were used as input. The cross-area dendrogram was generated with a similar workflow but was downsampled to a maximum of 100 nuclei per cross-area cluster per area. The 3000 High-variance genes used for dendrogram construction were identified from the downsampled matrix containing Cv3 nuclei from all eight areas.

### Cell type comparisons across cortical areas

#### Differential gene expression

To identify subclass marker genes within an area, Cv3 datasets from each area were downsampled to a maximum of 100 nuclei per cluster per individual. Differentially expressed marker genes were then identified using the FindAllMarkers function from Seurat, using the Wilcoxon sum rank test on log-normalized matrices with a maximum of 500 nuclei per group (subclass vs. all other nuclei as background). Statistical thresholds for markers are indicated in their respective figures. To identify area marker genes across subclasses, Cv3 datasets from each area were downsampled to a maximum of 50 nuclei per cluster per individual. Downsampled counts matrices were then grouped into pseudo-bulk replicates (area, individual, subclass) and the counts were summed per replicate. DESeq2 functionality was then used to perform a differential expression analysis between area pairs (or comparisons of interest) for each subclass using the Wald test statistic.

#### Transcriptomic entropy across areas

To quantify inter-cell transcriptomic heterogeneity across areas for each subclass we calculated the transcriptomic entropy in the observed data (structured) and compared against entropy in permuted data (unstructured). Transcriptomic heterogeneity is defined as the difference between the structured and unstructured entropy. To compute transcriptomic entropy we followed these steps: (1) Randomly down-sample the cells within each subclass by taking 250 cells from each cross area cell type. (2) Identify the highly variable genes in each area and take the union of genes as our set of interest. (3) Then, by following a recently reported computational approach to quantify transcriptomic heterogeneity (*72*), we computed the per-area transcriptomic entropy for each subclass.

#### Identifying changes in cell type proportions across areas

Cell type proportions are compositional, where the gain or loss of one population necessarily affects the proportions of the others, so we used scCODA (*73*) to determine which changes in cell class, subclass, and cell type proportions across areas were statistically significant. We analyzed neuronal and non-neuronal populations separately since nuclei were sorted based on NeuN immunostaining to enrich for neurons. The proportion of each cell type was estimated using a Bayesian approach where proportion differences across individuals were used to estimate the posterior. All compositional and categorical analyses require a reference population to describe differences with respect to and, because we were uncertain which populations should be unchanged, we iteratively used each cell type and each area as a reference when computing abundance changes. To account for sex differences, we included it as a covariate when testing for abundance changes. Separately for neuronal and non-neuronal populations, we reported the effect size of each area for each cell type (**Table S9**) and used a mean inclusion probability cutoff of 0.7 for calling a population consistently different.

#### Partitioning variation in gene expression across areas

Variation partitioning analysis was performed to prioritize the drivers of variation across areas within each subclass. Using linear mixed-effect models implemented in the variancePartitioning bioconductor package: http://bioconductor.org/packages/variancePartition (*74*) we identify genes whose variance is best explained along the mediolateral, rostrocaudal and dorsoventral axes as well as by cortical area and donor. The order of areas along these axes was defined based on the approximate x, y, and z coordinates of tissue samples based on a common coordinate framework of the adult human brain (*28*) (**Table S2**). Genes were removed from the analysis based on the following criteria: (1) expressed in less than 10 cells, (2) greater than 80% dropout rate, (3) zero variance in expression, and (4) expression less than 1 CPM on average. The variance partitioning linear mixed-effect model was then defined as:

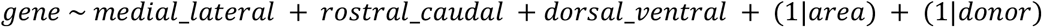

and passed into the variancePartition function ‘fitVarPartModel()’. We determined the amount of variation explained per covariate for each gene from the ‘extractVarPart()’ function.

### *In situ* profiling of gene expression

Human postmortem frozen brain tissue was embedded in Optimum Cutting Temperature medium (VWR,25608-930) and sectioned on a Leica cryostat at −17 C at 10 um onto Vizgen MERSCOPE coverslips (VIZGEN 2040003). These sections were then processed for MERSCOPE imaging according to the manufacturer’s instructions. Briefly: sections were allowed to adhere to these coverslips at room temperature for 10 min prior to a 1 min wash in nuclease-free phosphate buffered saline (PBS) and fixation for 15 min in 4% paraformaldehyde in PBS. Fixation was followed by 3×5 minute washes in PBS prior to a 1 min wash in 70% ethanol. Fixed sections were then stored in 70% ethanol at 4 C prior to use and for up to one month. Human sections were photobleached using a 150W LED array for 72 h at 4 C prior to hybridization then washed in 5 ml Sample Prep Wash Buffer (VIZGEN 20300001) in a 5 cm petri dish. Sections were then incubated in 5 ml Formamide Wash Buffer (VIZGEN 20300002) at 37 C for 30 min. Sections were hybridized by placing 50 ul of VIZGEN-supplied Gene Panel Mix onto the section, covering with parafilm and incubating at 37 C for 36-48 h in a humidified hybridization oven.

Following hybridization, sections were washed twice in 5 ml Formamide Wash Buffer for 30 min at 47 C. Sections were then embedded in acrylamide by polymerizing VIZGEN Embedding Premix (VIZGEN 20300004) according to the manufacturer’s instructions. Sections were embedded by inverting sections onto 110 ul of Embedding Premix and 10% Ammonium Persulfate (Sigma A3678) and TEMED (BioRad 161-0800) solution applied to a Gel Slick (Lonza 50640) treated 2×3 glass slide. The coverslips were pressed gently onto the acrylamide solution and allowed to polymerize for 1.5 h. Following embedding, sections were cleared for 24-48 h with a mixture of VIZGEN Clearing Solution (VIZGEN 20300003) and Proteinase K (New England Biolabs P8107S) according to the Manufacturer’s instructions. Following clearing, sections were washed twice for 5 min in Sample Prep Wash Buffer (PN 20300001). VIZGEN DAPI and PolyT Stain (PN 20300021) was applied to each section for 15 min followed by a 10 min wash in Formamide Wash Buffer. Formamide Wash Buffer was removed and replaced with Sample Prep Wash Buffer during MERSCOPE set up. 100 ul of RNAse Inhibitor (New England BioLabs M0314L) was added to 250 ul of Imaging Buffer Activator (PN 203000015) and this mixture was added via the cartridge activation port to a pre-thawed and mixed MERSCOPE Imaging cartridge (VIZGEN PN1040004). 15 ml mineral oil (Millipore-Sigma m5904-6X500ML) was added to the activation port and the MERSCOPE fluidics system was primed according to VIZGEN instructions. The flow chamber was assembled with the hybridized and cleared section coverslip according to VIZGEN specifications and the imaging session was initiated after collection of a 10X mosaic DAPI image and selection of the imaging area. For specimens that passed the minimum count threshold, imaging was initiated and processing completed according to VIZGEN proprietary protocol. Following processing and segmentation via MERSCOPE software, cells with fewer than 50 counts, or with an area outside the 100-300 um2 range were eliminated from the mapping process.

The 140 gene human cortical panel was selected using a combination of manual and algorithmic based strategies requiring a reference single cell/nucleus RNA-seq data set from the same tissue, in this case the human MTG snRNAseq dataset and resulting taxonomy (*11*). First, an initial set of High-confidence marker genes are selected through a combination of literature search and analysis of the reference data. These genes are used as input for a greedy algorithm (detailed below). Second, the reference RNA-seq data set is filtered to only include genes compatible with mFISH. Retained genes need to be 1) long enough to allow probe design (> 960 base pairs); 2) expressed highly enough to be detected (FPKM >= 10), but not so high as to overcrowd the signal of other genes in a cell (FPKM < 500); 3) expressed with low expression in off-target cells (FPKM < 50 in non-neuronal cells); and 4) differentially expressed between cell types (top 500 remaining genes by marker score20). To more evenly sample each cell type, the reference data set is also filtered to include a maximum of 50 cells per cluster.

The main step of gene selection uses a greedy algorithm to iteratively add genes to the initial set. To do this, each cell in the filtered reference data set is mapped to a cell type by taking the Pearson correlation of its expression levels with each cluster median using the initial gene set of size n, and the cluster corresponding to the maximum value is defined as the “mapped cluster”. The “mapping distance” is then defined as the average cluster distance between the mapped cluster and the originally assigned cluster for each cell. In this case a weighted cluster distance, defined as one minus the Pearson correlation between cluster medians calculated across all filtered genes, is used to penalize cases where cells are mapped to very different types, but an unweighted distance, defined as the fraction of cells that do not map to their assigned cluster, could also be used. This mapping step is repeated for every possible n+1 gene set in the filtered reference data set, and the set with minimum cluster distance is retained as the new gene set. These steps are repeated using the new get set (of size n+1) until a gene panel of the desired size is attained. Code for reproducing this gene selection strategy is available as part of the mfishtools R library (https://github.com/AllenInstitute/mfishtools).

#### Cell type mapping of MERSCOPE data

Any genes not matched across both the MERSCOPE gene panel and the snRNASeq mapping taxonomy were filtered from the snRNASeq dataset. We calculated the mean gene expression for each gene in each snRNAseq cluster. We assigned MERSCOPE cells to snRNAseq clusters by finding the nearest cluster to the mean expression vectors of the snRNASeq clusters using the cosine distance. All scripts and data used are available at: https://github.com/AllenInstitute/human_cross_areal.

### GFAP Immunofluorescence

Tissue blocks from cortical areas of interest were removed from fresh-frozen tissue slabs as described above. Immediately after dissection, tissue blocks were drop-fixed in cold 4% paraformaldehyde overnight in a 4°C fridge. Tissue blocks were then rinsed in multiple washes of 1X PBS, cryoprotected in sequential 15% and 30% sucrose solutions, and embedded in OCT. Sections were cut free floating at 30μm in the coronal plane on a Leica cryostat into 1X PBS and were stored at 4°C or at −20°C in cryoprotectant solution until the time of use. Sections were processed for immunofluorescence using a rabbit polyclonal anti-GFAP antibody (Agilent, Z0334) at a dilution of 1:1000 and mouse monoclonal anti-NeuN antibody (Millipore Sigma, MAB377) at a dilution of 1:1000. Primary antibodies were incubated overnight at 4°C, followed by incubation in Alexa Fluor conjugated secondary species-specific antibodies for 2 hours at room temperature. Sections were counterstained with DAPI and Neurotrace 500 fluorescent Nissl stain and were mounted in ProLong Gold Antifade Mountant (ThermoFisher Scientific). Sections were imaged on a Nikon TiE fluorescence microscope equipped with NIS-Elements Advanced Research imaging software (v4.20). GFAP processes were traced using the SNT plugin in the Fiji distribution of ImageJ.

## Acknowledgements

We thank the tissue procurement, tissue processing and facilities teams at the Allen Institute for Brain Science for assistance with the transport and processing of postmortem and neurosurgical brain specimens; the technology team at the Allen Institute for assistance with data management; M. Vawter, J. Davis and the San Diego Medical Examiner’s Office for assistance with postmortem tissue donations. Research reported in this publication was supported by the National Institute Of Mental Health of the National Institutes of Health under Award Numbers U01MH114812 and U19MH114830. The content is solely the responsibility of the authors and does not necessarily represent the official views of the National Institutes of Health. The authors thank the founder of the Allen Institute, Paul G. Allen, for his vision, encouragement and support.

## Author contributions

RNA data generation: AMY, AT, BPL, CDK, CR, DB, DH, DM, ERB, ESL, HT, JG, JS, KC, KL, KSi, KSm, KW, MKr, MT, ND, NS, RDH, SD, SL, SSh, TC, TEB, TP
Spatial transcriptomic data generation: AR, BL, DM, EG, JCa, JCl, MKu, NM
Data archive / Infrastructure: JG, SSo
Data analysis: BL, EG, ESL, JCl, JG, KJT, KSm, NJ, NLJ, TEB
Data interpretation: EMC, ESL, JCl, JG, KJT, NJ, NLJ, PPM, PRH, RDH, SD, TEB
Writing manuscript: ESL, JCl, KJT, NLJ, PRH, RDH, TEB

## Data and materials availability

Raw sequence data were produced as part of the BRAIN Initiative Cell Census Network (BICCN: RRID:SCR_015820) are available for download from the Neuroscience Multi-omics Archive (RRID:SCR_016152; https://assets.nemoarchive.orq/dat-rq2rc5m) and the Brain Cell Data Center (RRID:SCR_017266; https://biccn.org/data). Code for analysis and generation of figures is available for download from https://github.com/AllenInstitute/human_cross_areal. MTG human SMARTseq v4 data (https://portal.brain-map.org/atlases-and-data/rnaseq/human-mtg-smart-seq, https://assets.nemoarchive.org/dat-swzf4kc).

## Supplementary Materials

Figs. 1 to 12

Tables 1 to 13 legends

