## Supplemental Figures 1-12 and Table legends 1-13 for "Transcriptomic cytoarchitecture reveals principles of human neocortex organization"

### Transcriptomic specialization of functional areas in human neocortex

**This PDF file includes:**

Figs. S1 to S12
Tables S1 to S13 legends

### *Figure S1*

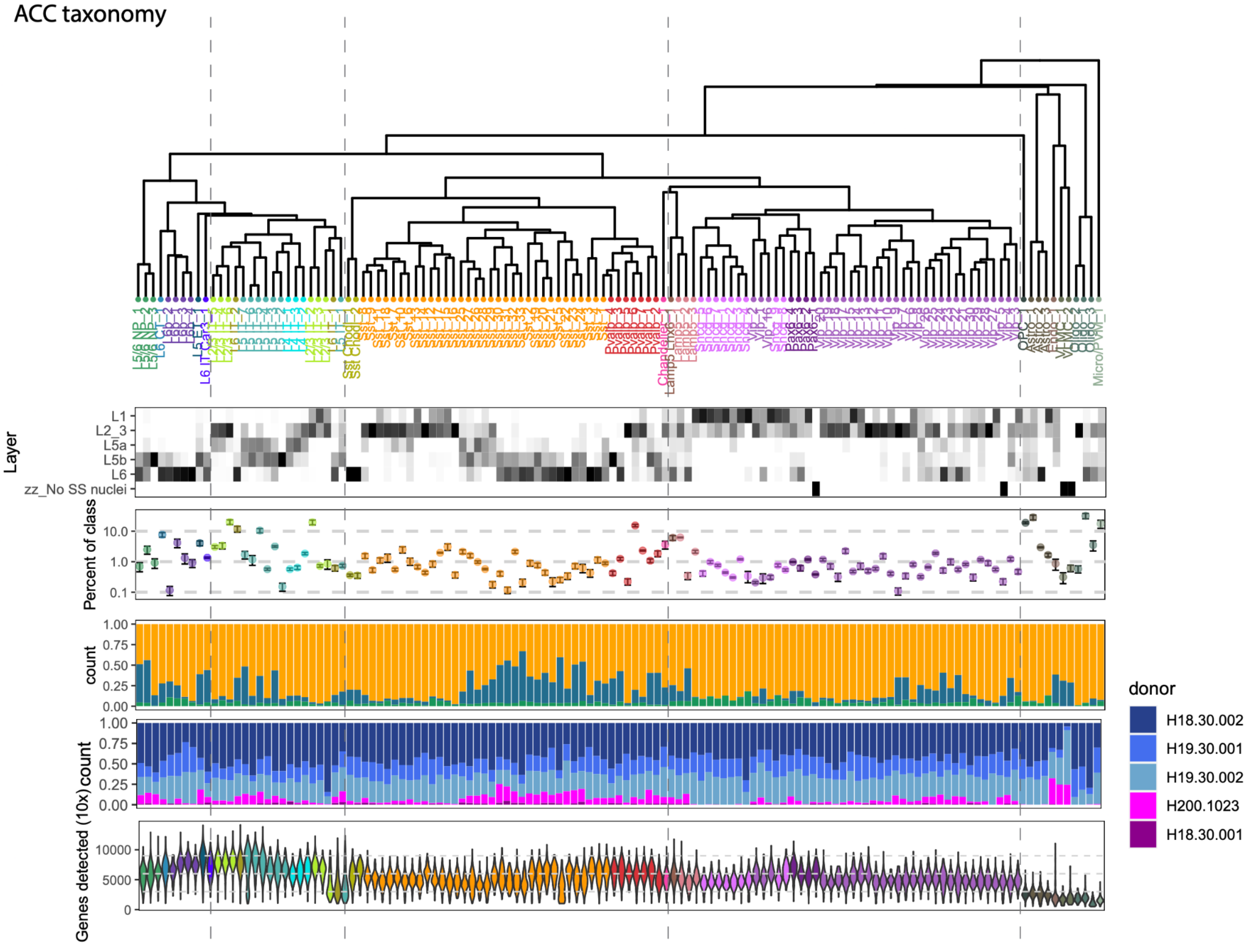


**Fig. S1. Anterior cingulate cortex (ACC) taxonomy and cell type features.** Taxonomy of cell types, laminar distributions estimated based on SSv4 layer dissections, proportions of cell class, relative sampling by dataset (Cv3, orange; Cv3 layer 5, blue; SSv4, green) and donor, and genes detected.

### *Figure S2*

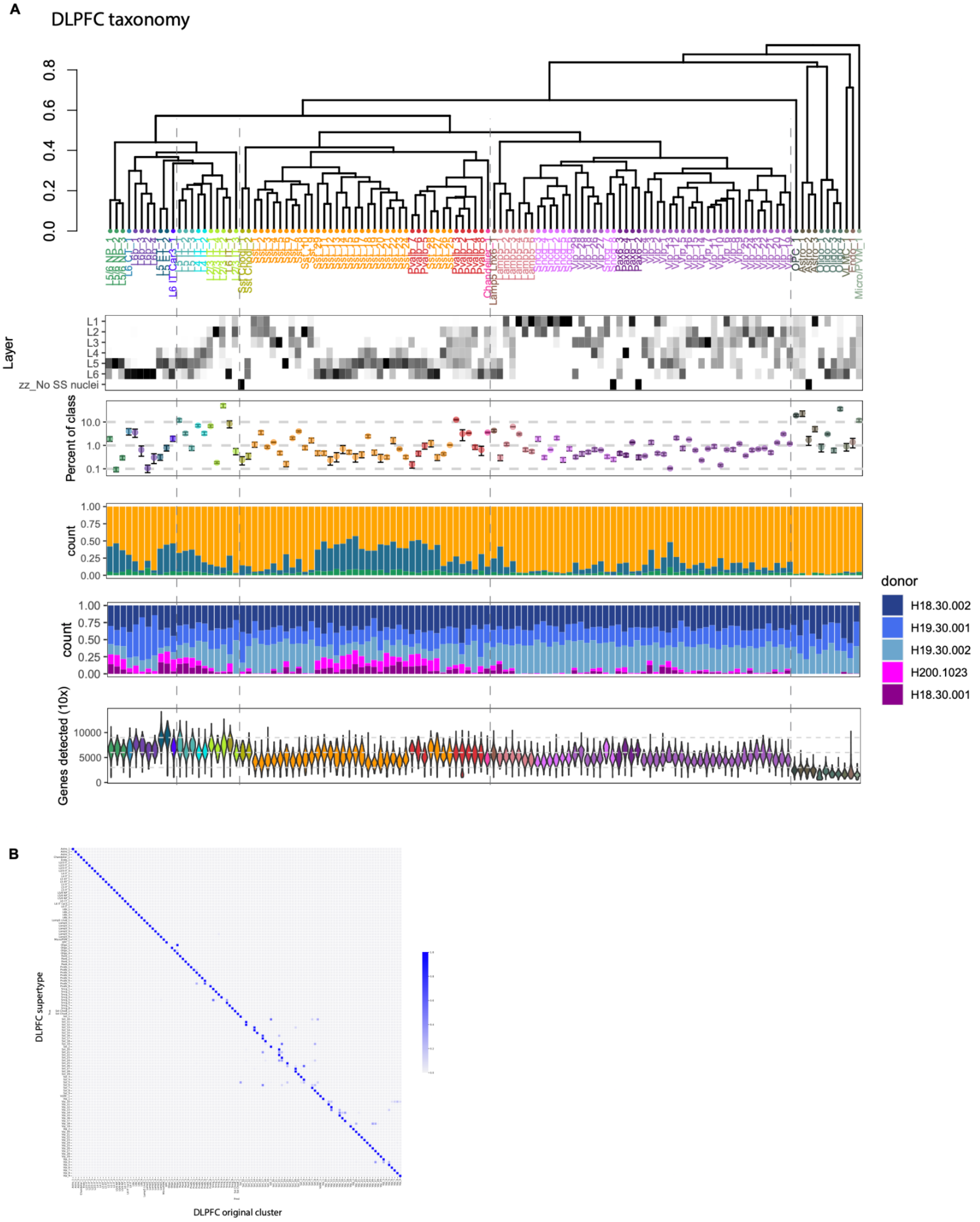


**Fig. S2. Dorsolateral prefrontal cortex (DFC) taxonomy and cell type features. (A)** Taxonomy and features of cell types. **(B)** Confusion matrix showing the mapping of robustly mappable supertypes (DFC) to the taxonomy clusters. Note that most supertypes map 1-to-1 to a cluster, while a few supertypes map to several highly similar clusters that could not be robustly distinguished.

*Figure S3*

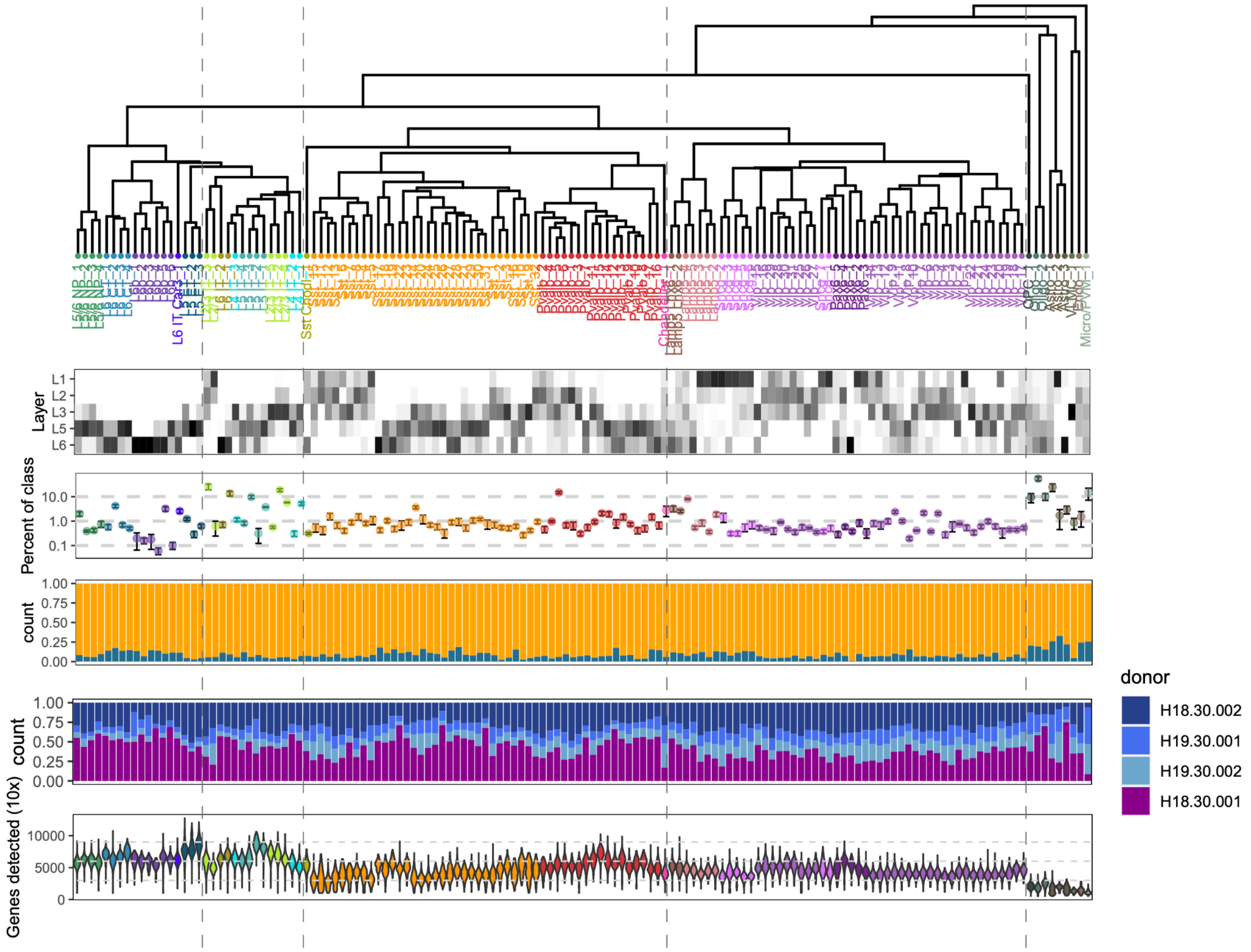


**Fig. S3. Primary motor cortex (M1) taxonomy and cell type features.**

*Figure S4*

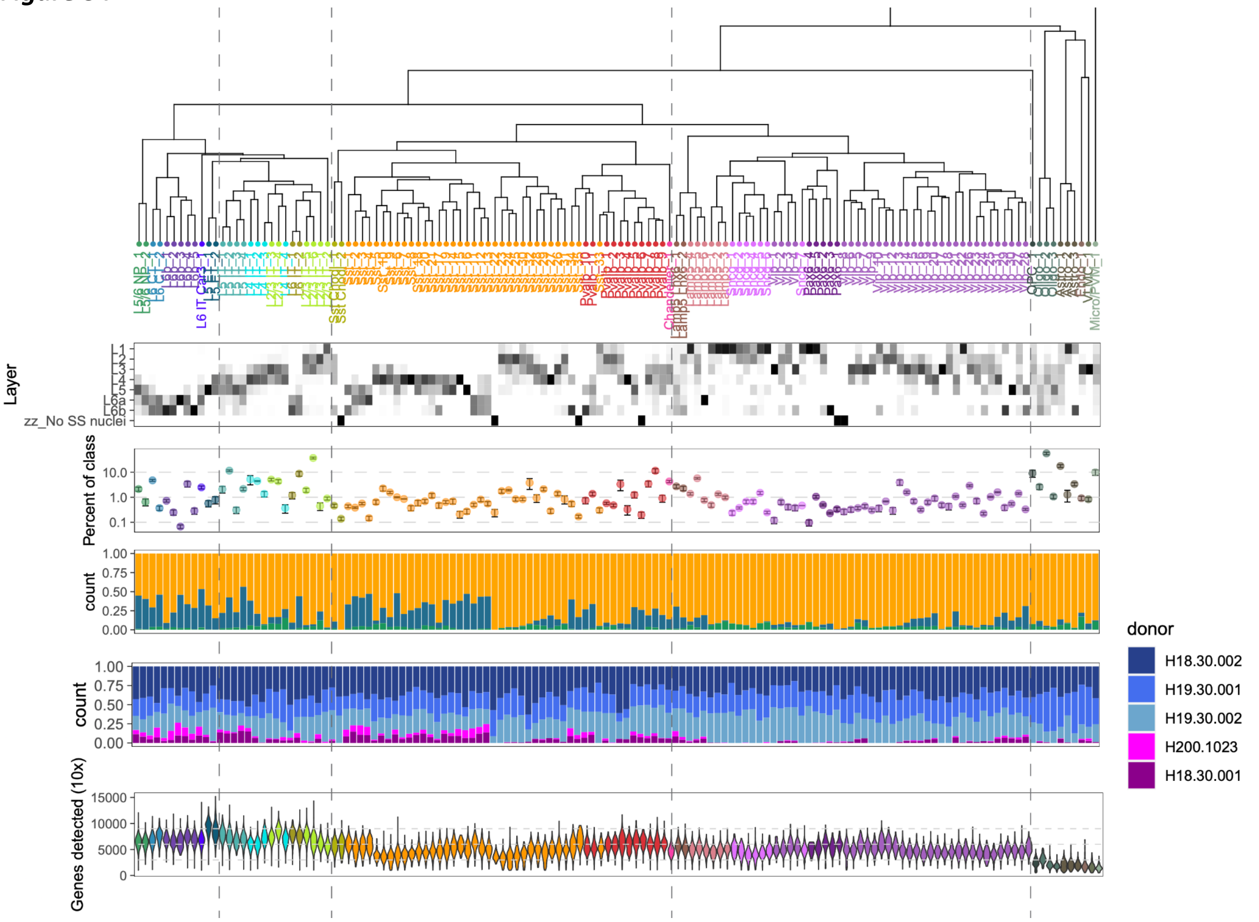


**Fig. S4. Primary somatosensory cortex (S1) taxonomy and cell type features.**

*Figure S5*

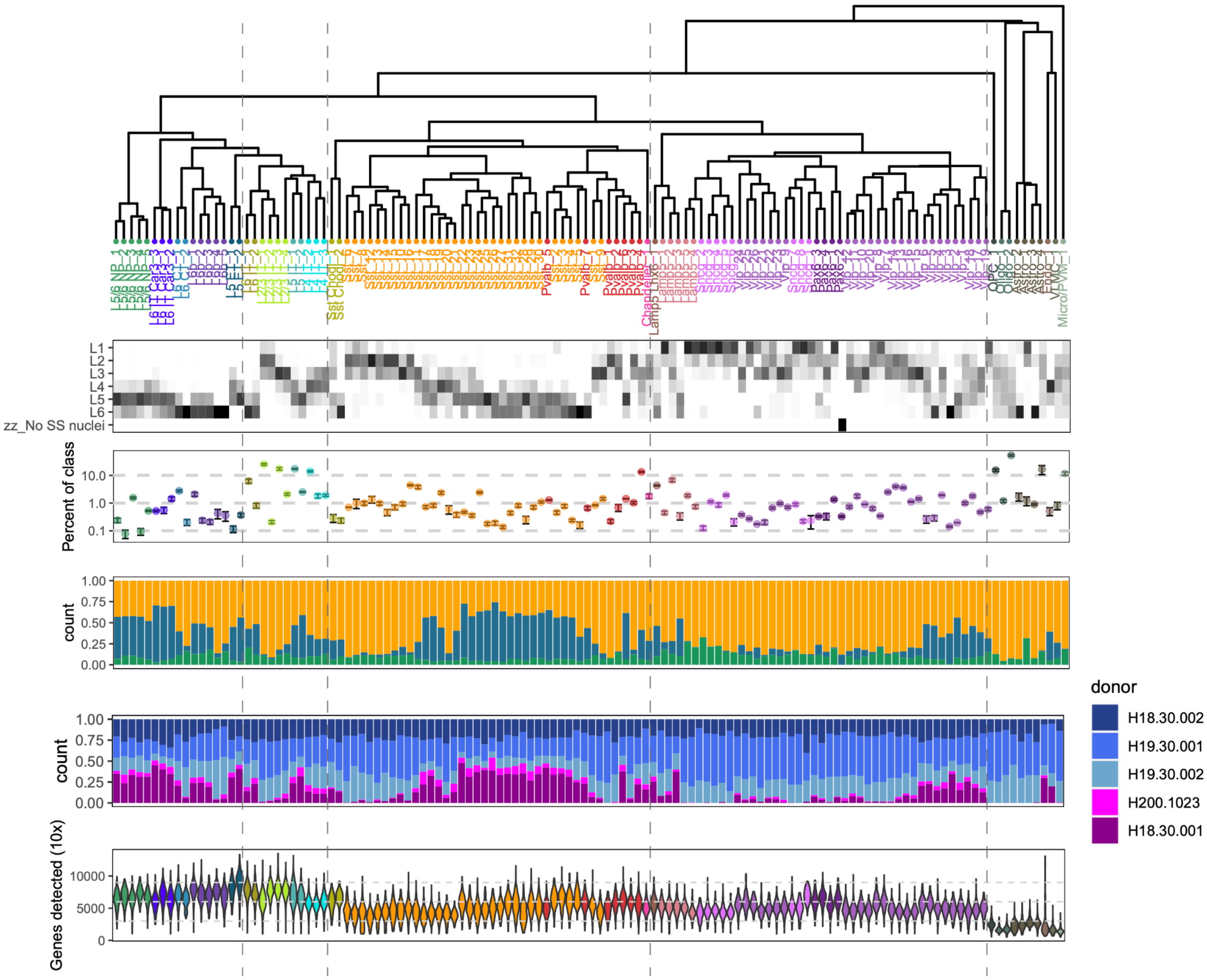


**Fig. S5. Middle temporal gyrus (MTG) taxonomy and cell type features.**

### *Figure S6*

A1 taxonomy

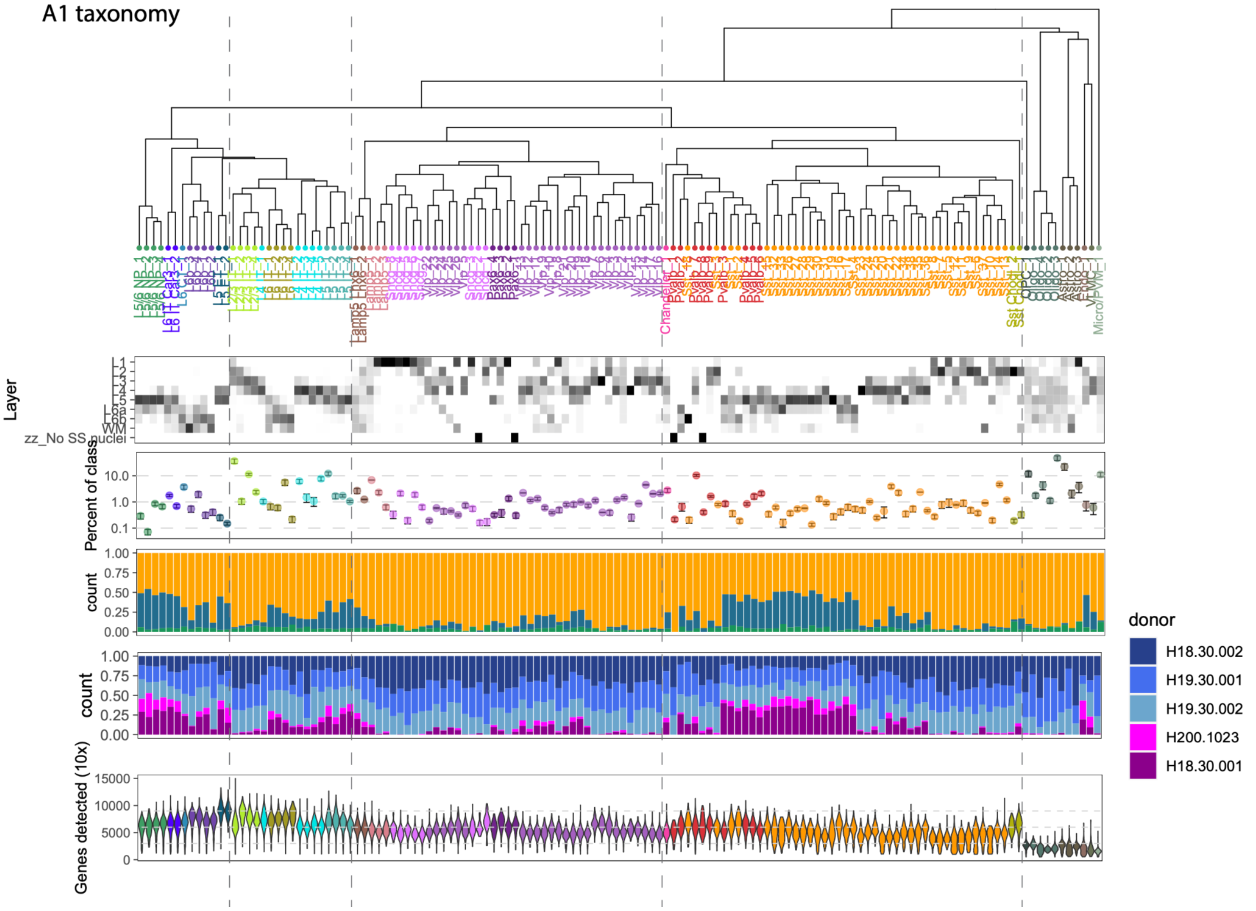


**Fig. S6. Primary auditory cortex (A1) taxonomy and cell type features.**

*Figure S7*

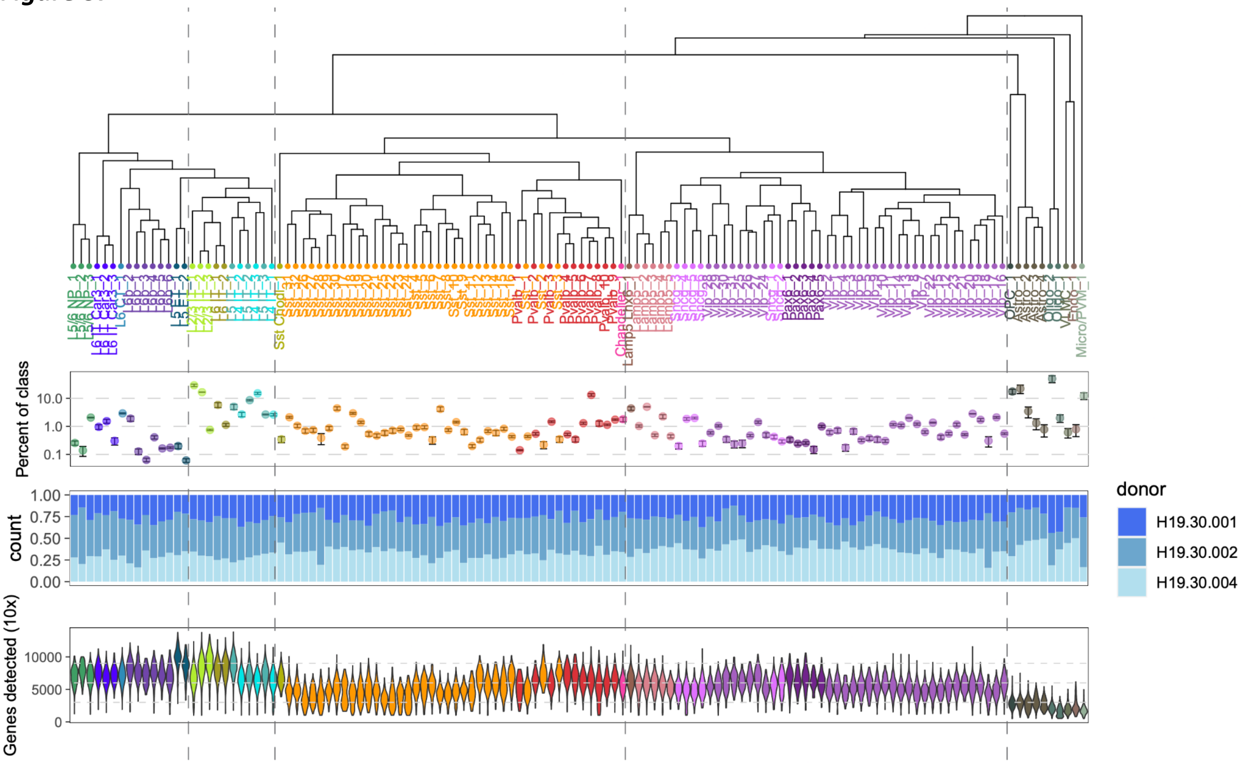


**Fig. S7. Angular gyrus (AnG) taxonomy and cell type features.**

***Figure S8***

V1 taxonomy

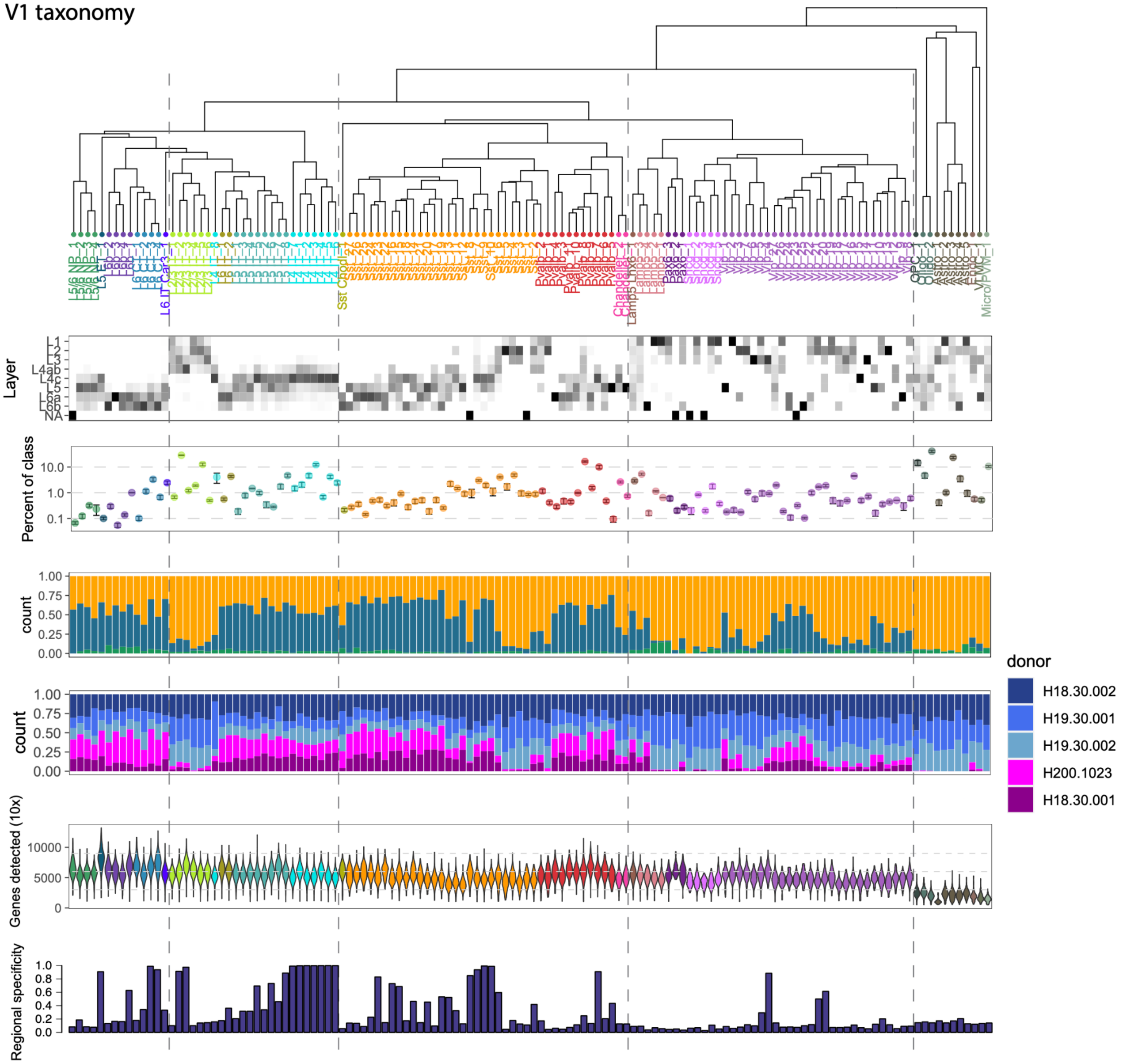


**Fig. S8. Primary visual cortex (V1) taxonomy and cell type features.** Areal specificity quantifies the transcriptomic distinctiveness of clusters in V1 compared to other areas. Each V1 cluster maps to one or more cross–area consensus types, as reported in **Table S11**. The specificity was calculated as the sum of the products of the proportion of V1 nuclei that map to each consensus type times the proportion of nuclei from V1 in that consensus type.

### *Figure S9*

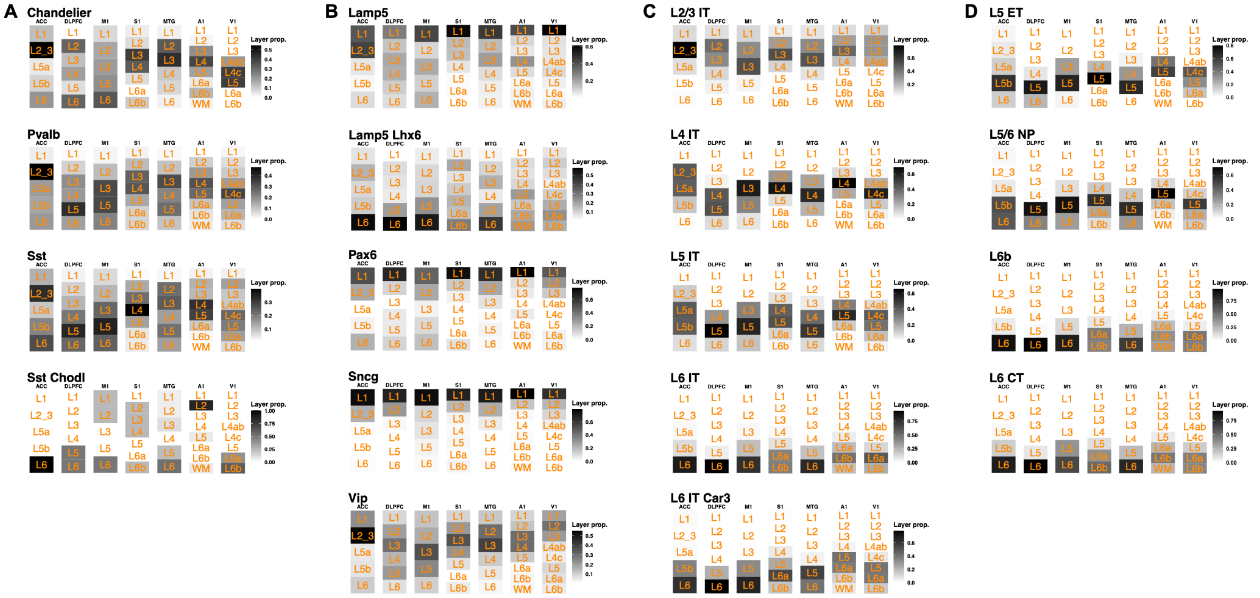


**Fig. S9. Laminar distributions of neuronal subclasses across areas. (A-D)** Estimated laminar distributions based on layer dissections from seven cortical areas for MGE-derived (A) and CGE-derived (B) interneuron subclasses and IT-projecting (C) and deep layer, non-IT-projecting (D) excitatory neuron subclasses.

***Figure S10***

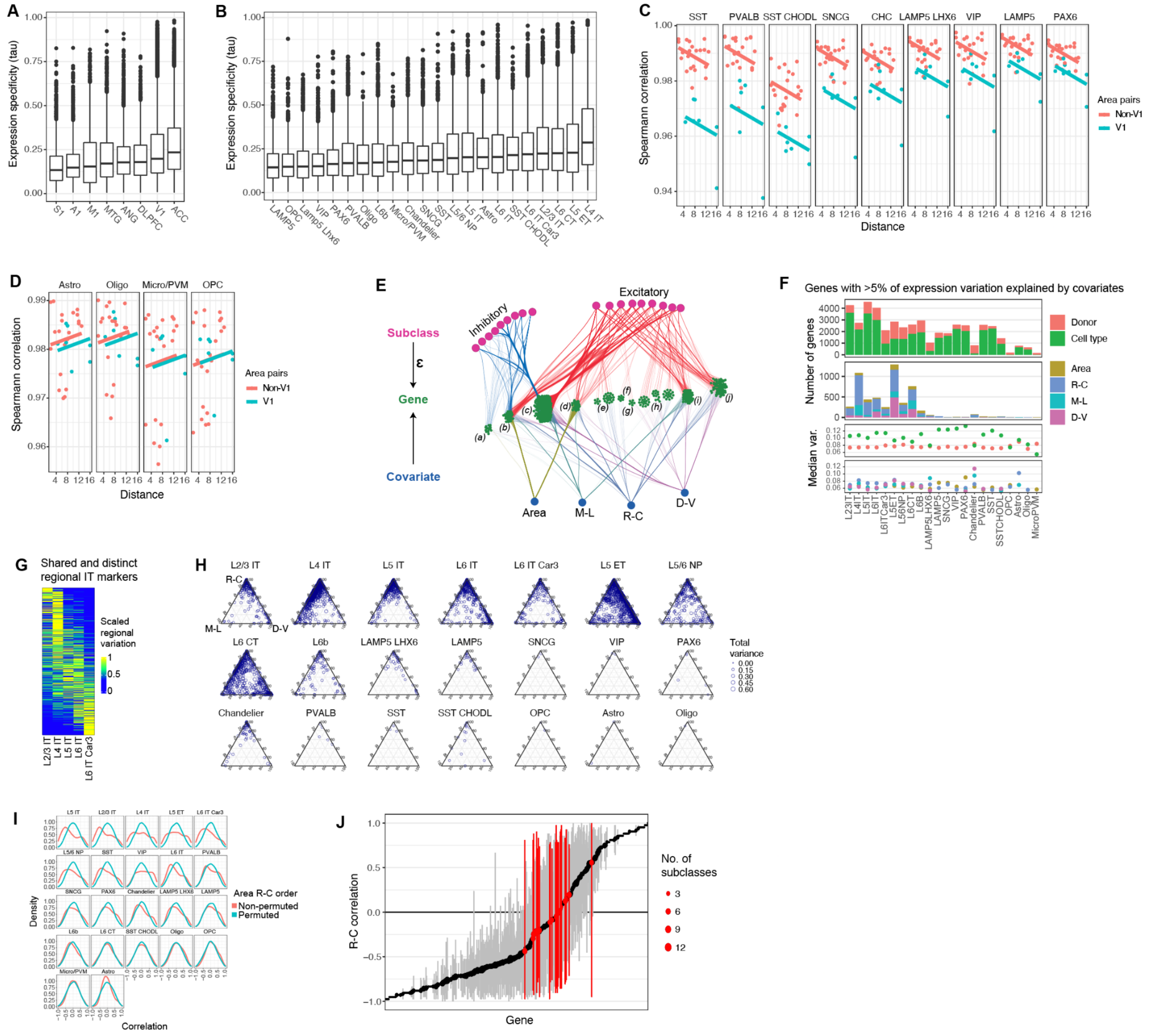


**Fig. S10. Transcriptomic topography of adult human cortex. (A)** Distributions of subclass expression specificity grouped by area with maximum expression. ACC and V1 have the most distinct transcriptomic signatures, and S1 and A1 have the least. **(B)** Distributions of subclass expression specificity grouped by subclass and with maximum expression in any area. **(C,D)** Expression similarity of inhibitory neuron types (C) and non-neuronal types (D) between pairs of areas that include V1 (blue) or do not include 1 V1 (red) as a function of physical distance (cm) on the cortical sheet. **(E)** Graph summarizing the results of a variance partitioning analysis showing all genes with >5% of variation in expression explained by an areal covariate. Lines connect areal covariates and inhibitory (blue) and excitatory (red) neuron subclasses to genes if >5% of the variance is explained. Genes are grouped based on shared connectivity in the graph and labels correspond to gene lists in **Table S8**. Note that genes in group (a) have areal patterning restricted to inhibitory subclasses despite much stronger gradients among excitatory neurons overall. **(F)** Summary of the number of genes with >5% of variation in gene expression explained by covariates in a variation partitioning analysis. Areal patterning and other covariates are separated into two plots with different scales of gene counts. The cell type covariate represents clusters from within-area taxonomies.

**(G)** Heatmap of the relative proportions of expression variance explained by region for IT neuron subclasses. Each row corresponds to a gene, and rows sum to 1. **(H)** Ternary plots summarizing the relative proportion of expression variance explained by the position of regions along R-C, M-L, and D-V gradients for each subclass. All genes are shown that have >5% expression variance explained by at least one gradient, and the point size indicates the total variance explained by the gradients (5-60%). **(I)** Distributions of correlations of subclass expression with areas ordered by rostrocaudal cortical position and with areas ordered randomly. Permuted correlations are approximately normally distributed, while non-permuted correlations often include higher and lower values. This supports the finding that R-C and P-A expression gradients are stronger than would be expected by chance. **(J)** For genes with an R-C gradient in at least one subclass (shown in Fig. 2F), the range of R-C correlations is plotted for all subclasses with significantly variable expression across areas. Genes are ordered by the median correlation, and the number of subclasses included in the range are indicated by point size. The minority of genes that have opposing expression gradients in different subclasses (i.e. correlations < -0.7 and > 0.7 are included in the range) are highlighted in red and can be identified in **Table S7**.

*Figure S11*

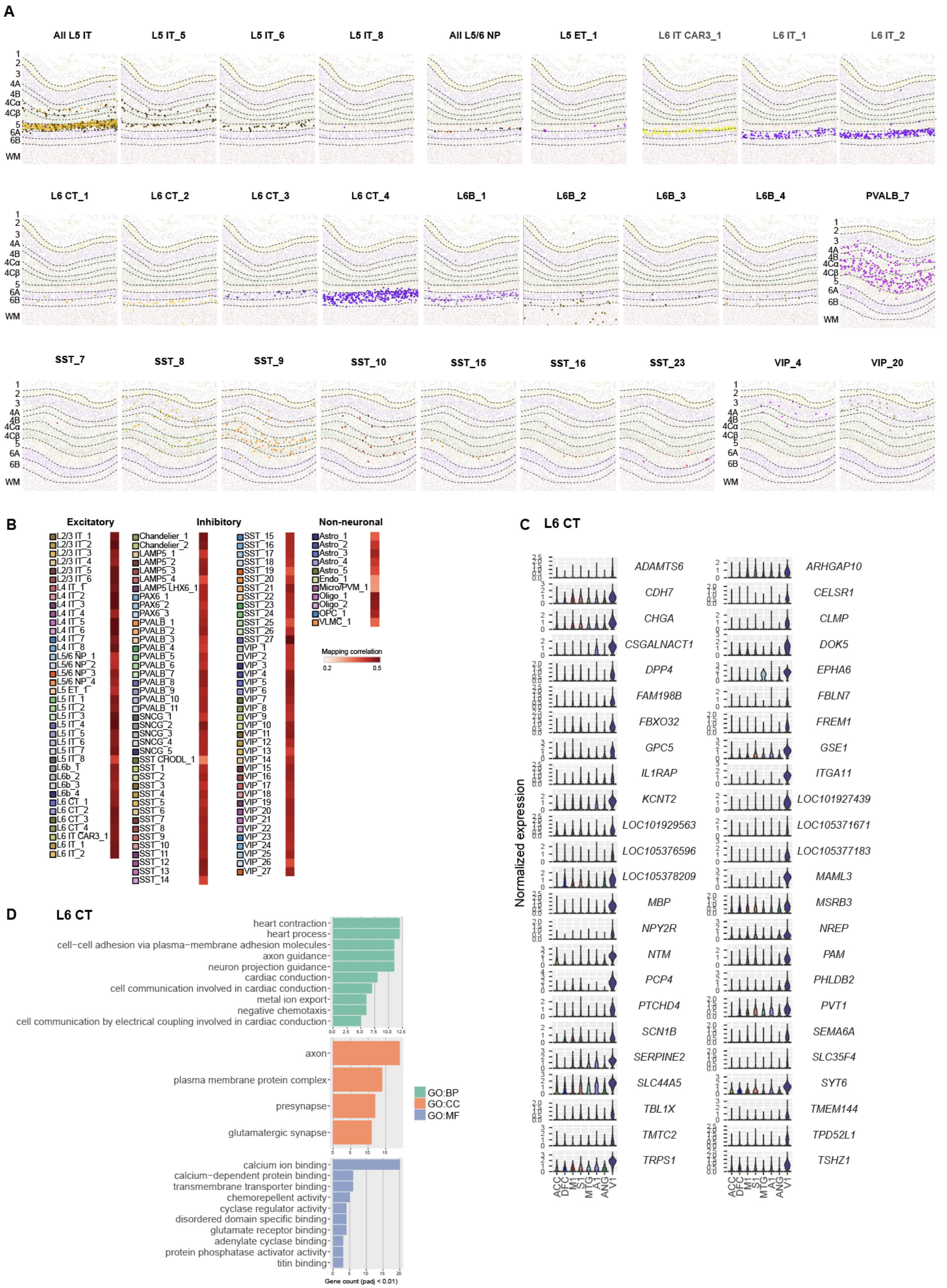

A
All L5 IT
L5 IT_5
L5 IT_6
L5 IT_8
All L5/6 NP
L5 ET_1
L6 IT CAR3_1
L6 IT_1
L6 IT_2
L6 CT_1
L6 CT_2
L6 CT_3
L6 CT_4
L6B_1
L6B_2
L6B_3
L6B_4
PVALB_7
SST_7
SST_8
SST_9
SST_10
SST_15
SST_16
SST_23
VIP_4
VIP_20
1
2
3
4A
4B
4Cα
4Cβ
5
6A
6B
WM
B
Excitatory
Inhibitory
Non-neuronal
Mapping correlation
0.2
0.5
C
L6 CT
Normalized expression
ADAMTS6
ARHGAP10
CDH7
CELSR1
CHGA
CLMP
CSGALNACT1
DOK5
DPP4
EPHA6
FAM198B
FBLN7
FBXO32
FREM1
GPC5
GSE1
IL1RAP
ITGA11
KCNT2
LOC101927439
LOC101929563
LOC105371671
LOC105376596
LOC105377183
LOC105378209
MAML3
MBP
MSRB3
NPY2R
NREP
NTM
PAM
PCP4
PHLDB2
PTCHD4
PVT1
SCN1B
SEMA6A
SERPINE2
SLC35F4
SLC44A5
SYT6
TBL1X
TMEM144
TMTC2
TPD52L1
TRPS1
TSHZ1
ACC
DFC
M1
S1
MTG
A1
ANG
V1
D
L6 CT
heart contraction
heart process
cell−cell adhesion via plasma−membrane adhesion molecules
axon guidance
neuron projection guidance
cardiac conduction
cell communication involved in cardiac conduction
metal ion export
negative chemotaxis
cell communication by electrical coupling involved in cardiac conduction
axon
plasma membrane protein complex
presynapse
glutamatergic synapse
calcium ion binding
calcium−dependent protein binding
transmembrane transporter binding
chemorepellent activity
cyclase regulator activity
disordered domain specific binding
glutamate receptor binding
adenylate cyclase binding
protein phosphatase activator activity
titin binding
GO:BP
GO:CC
GO:MF
Gene count (padj < 0.01)

**Fig. S11. Spatial distributions of V1 cell types. (A)** Laminar distributions of cell types based on MERFISH *in situ* labeling experiments of human V1 tissue. **(B)** Mapping confidence of cells to reference V1 types based on correlated expression of marker genes. Most cell types have similar mapping strength, with reduced correlations for SST CHODL, endothelial cells, and microglia likely due to fewer marker genes being included in the MERFISH gene panel. **(C)** Regional markers of L6 CT neurons that have selectively enriched expression in V1. **(D)** GO enrichment analysis of V1 markers of L6 CT neurons.

### *Figure S12*

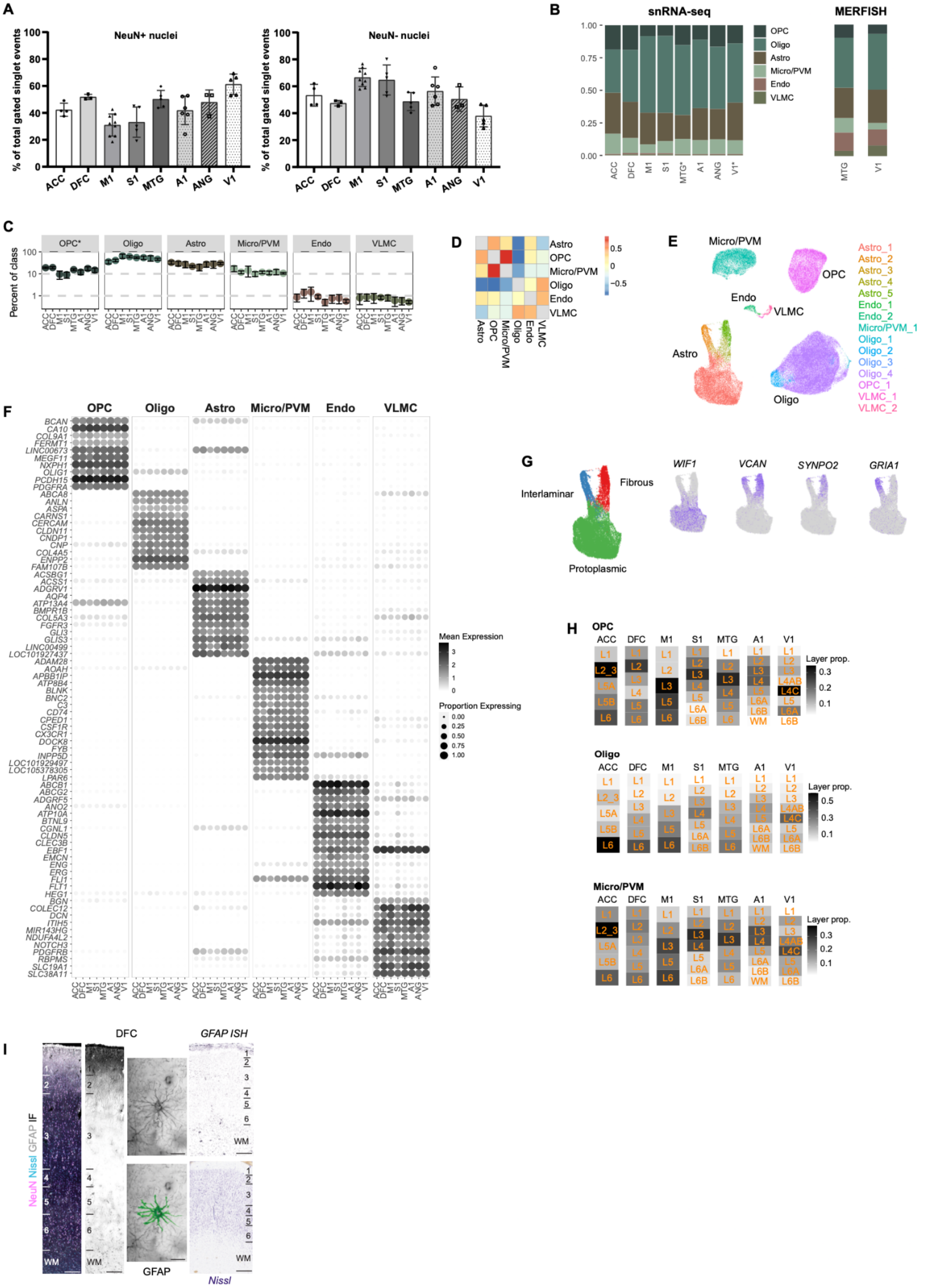
A
NeuN+ nuclei
NeuN- nuclei
% of total gated singlet events
ACC DFC M1 S1 MTG A1 ANG V1
B
snRNA-seq
MERFISH
OPC
Oligo
Astro
Micro/PVM
Endo
VLMC
C
Percent of class
D
E
Astro_1
Astro_2
Astro_3
Astro_4
Astro_5
Endo_1
Endo_2
Micro/PVM_1
Oligo_1
Oligo_2
Oligo_3
Oligo_4
OPC_1
VLMC_1
VLMC_2
F
Mean Expression
Proportion Expressing
G
Interlaminar
Fibrous
Protoplasmic
WIF1
VCAN
SYNPO2
GRIA1
H
Layer prop.
I
DFC
GFAP ISH
NeuN Nissl GFAP IF
GFAP
Nissl

**Fig. S12. Non-neuronal diversity across areas. (A)** Bar plots comparing the percentage of nuclei from dissociated tissue that were NeuN+ (neuronal) or NeuN- (non-neuronal) from each cortical area assessed using flow cytometry. Bars show the mean (bar height), error bars the standard deviation, and points represent values for individual replicates. **(B)** Relative proportions of non-neuronal subclasses as a fraction of all non-neuronal cells in each area and estimated based on snRNA-seq profiling or *in situ* labeling using MERFISH. The tissue dissociation protocol was not optimized to capture vascular cells, and therefore endothelial cells and VLMCs were underrepresented in the snRNA-seq data. **(C)** Non-neuronal subclass proportions across areas. Significant differences across regions (ANOVA; *nominal $P < 0.05$). **(D)** Spearman correlations of non-neuronal subclass proportions across areas. **(E)** UMAP of glial cells labeled by cross-area consensus type. **(F)** Conserved markers of non-neuronal subclasses across areas (see related **Table S4** and **S9**). **(G)** Conserved markers of astrocyte subtype across areas. **(H)** Estimated laminar distributions based on layer dissections from seven cortical areas for glial subclasses. **(I)** GFAP immunofluorescence (IF) and *in situ* hybridization (ISH) illustrates variable laminar distributions and morphologies of astrocytes in DFC. Scale bars: IF columns (100µm), GFAP tracing images (15 µm), ISH (200 µm).

*Supplementary Tables*

**Table S1: Neuronal and non-neuronal subclass descriptions.**

**Table S2: Spatial locations of cortical areas.** Common coordinate framework (CCF) coordinates based on an adult human atlas (*28*). Also, approximate physical distances between areas on an unfolded neocortical sheet.

**Table S3: Cell type markers for within-area taxonomies.** Genes with significantly differential expression in each cell type versus all other cell types.

**Table S4: Subclass markers for within-area taxonomies.** Genes with significantly differential expression in each subclass versus all other subclasses.

**Table S5: Subclass proportions and E:I ratios across donors, areas, and layers.** Proportions and E:I ratios were estimated based on cell counts in MERFISH *in situ* experiments and based on snRNA-seq profiling of tissue dissection of the full depth of cortex (Cv3) or individual layers (SSv4).

**Table S6: Subclass DEGs across areas.** Genes with significantly differential expression across areas for each subclass based on ANOVA tests. Nominal P-values are reported that remain significant after Bonferroni correction for multiple testing of genes.

**Table S7: Genes with area-enriched and rostrocaudal gradient expression.**

**Table S8: Variance partitioning analysis results.**

**Table S9: Consensus cell type markers.** Markers of each consensus type versus all other types that are significantly differentially expressed (Bonferroni-adjusted P-value < 0.01, log2(fold-change) > 1) in at least one region. Nominally significant (P-value < 0.01) log2(fold-change) values are reported for all types and regions.

**Table S10: Compositional changes in consensus types across cortical areas.** Effect sizes of compositional changes in cell types across areas and between males and females were estimated using scCODA. Effect sizes were set to 0 if the mean inclusion probability was <0.7.

**Table S11: Area-specificity of within-area cell types.** Transcriptomic distinctiveness of cell types in each area compared to the most similar cell types in other areas. For example, for V1 the specificity was calculated as the sum of the products of the proportion of V1 nuclei that map to each consensus type times the proportion of nuclei from V1 in that consensus type. A score equal to 1 indicates that a cell type clusters completely separately from other areas in the consensus taxonomy, such as the V1-specific L4 IT types.

**Table S12: Markers of L2/3 IT, L4 IT, and SST types in V1.** Reported genes are significantly differentially expressed between each type and the corresponding subclass, for example, L2/3 IT_2 markers compared to all L2/3 IT neurons.

**Table S13: Genes with areal-enriched expression in L5 ET neurons.** DEGs that have significantly higher expression in L5 ET neurons in each area compared to all other areas.
